# Primary vitamin B_12_ limitation and widespread stress in Antarctic phytoplankton

**DOI:** 10.64898/2026.11.05.764676

**Authors:** Robert H. Lampe, Deepa Rao, Riss M. Kell, Rebecca J. Chmiel, Nicole L. Schanke, Soo Yoon Kim, Lauren E. Lees, Francesco Bolinesi, Olga Mangoni, Raffaella Casotti, Cecilia Balestra, Hong Zheng, Dawn M. Moran, Andrew E. Allen, Matthew R. McIlvin, Giacomo R. DiTullio, Mak A. Saito

## Abstract

Iron is considered the dominant limiting or colimiting nutrient for phytoplankton growth in the Southern Ocean. However, strong competition and limited biosynthesis capabilities for vitamin B_12_ (cobalamin) within the microbial community creates conditions suitable for cobalamin limitation. Through a nutrient enrichment experiment in the Amundsen Sea, we show that growth in a phytoplankton community comprising diatoms and prasinophytes was limited primarily by cobalamin and secondarily by iron. Protein biomarkers across a broad area of the Southern Ocean further indicate widespread cobalamin stress within diatoms and *Phaeocystis*. As warming-induced melting of ice-shelves increases iron supplies in the coastal Southern Ocean, shifts from iron to cobalamin limitation favor mixotrophic flagellates with low cobalamin requirements, altering community composition and function.

## Introduction

The coastal polynyas of the Southern Ocean are areas of open water surrounded by ice that disproportionally contribute to primary production and harbor high abundances of eukaryotic phytoplankton such as diatoms and the colonial haptophyte, *Phaeocystis antarctica*^1,2^. Other eukaryotic phytoplankton lineages, such as prasinophytes and cryptophytes, are also prominent, especially near the Western Antarctic Peninsula^3^. Together as primary producers, these phytoplankton communities play a critical role in supporting higher trophic levels and Earth’s biogeochemical cycles, including sequestering approximately 10% of carbon globally^4–7^.

Phytoplankton communities in the Southern Ocean are often limited by the micronutrient iron or light^8,9^. Other micronutrients, such as manganese, zinc, and vitamin B_12_ (cobalamin), have been shown to regulate growth of phytoplankton communities in the Southern Ocean, often as co-limiting nutrients with iron^10,11^. While manganese limitation may be restricted to open ocean regions^10,12^, cobalamin has been found to co-limit or be the second limiting nutrient with iron in coastal communities^13,14^. However, the extent of cobalamin limitation remains unconstrained.

In Antarctic polynyas, trace metal inputs are primarily related to physical processes such as upwelling, resuspension from margin sediments, and glacial or ice shelf melt^15–17^. Iron recycling through grazing further increases supply and enables blooms to persist^18,19^. In contrast, cobalamin is only biologically synthesized by certain prokaryotes, using the trace metal cobalt as a cofactor^20^. However, dissolved cobalt inventories in the Southern Ocean are often low (< 40 pmol L^-1^), with most strongly bound to organic ligands, and bioavailable inorganic cobalt is rapidly taken up by both prokaryotes and eukaryotes^21^. Once in seawater, cobalamin photodegrades within days or is consumed within hours, largely for methionine synthesis^13,20,22,23^. As a result, the Southern Ocean experiences low cobalamin inventories, with observations in coastal Antarctica ranging from 0.03-4.0 pmol L^-1^ ^24–26^.

To investigate cobalamin limitation in the Southern Ocean, an incubation experiment was performed during austral summer with a natural microbial community from the near surface of the central Amundsen Sea Polynya (ASP) overlying the Dotson-Getz Trough (Station 4) (Fig. 1a-c), one of the most biologically productive polynyas in the Southern Ocean^27^. Experimental treatments comprised iron and cobalamin addition, both separately (+Fe or +B_12_) and in combination (+FeB_12_). An additional treatment contained the strong metal chelator, desferrioxamine B (+DFB) which reduces iron bioavailability. Physiological and proteomic responses are evaluated both within the whole phytoplankton community and dominant groups alongside biogeochemical characterization of the experimental location. With metaproteomic indicators, we provide additional evidence of cobalamin stress in phytoplankton along a 10,000 km transect in the Southern Ocean.

**Figure 1.**
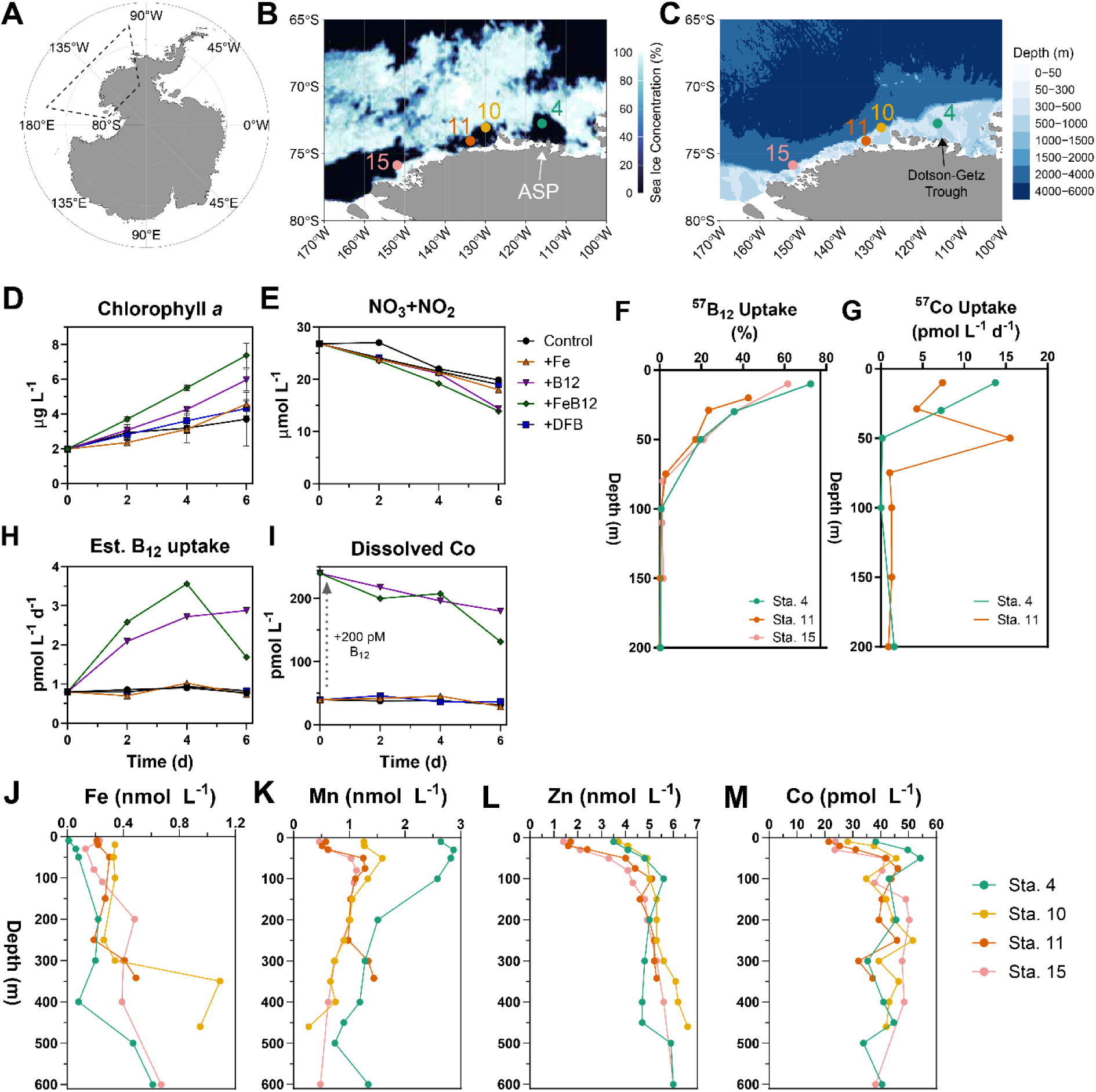
(A) Map of Antarctica with the study area outlined. (B-C) Sea ice concentration from the date of the experiment and bathymetry maps of the Amundsen Sea with stations denoted. (D) Chlorophyll *a* (µg L^-1^) and (E) nitrate+nitrite concentrations (µmol L^-1^) among the experimental treatments over time. (F-G) Vertical profiles of cobalamin uptake (%) and cobalt uptake rates (pmol L^-1^ d^-1^) at stations 4, 11, and 15. (H-I) Estimated cobalamin uptake rates (pmol L^-1^ d^-1^) and total dissolved cobalt concentrations (pmol L^-1^) among the experimental treatments over time. (J-M) Vertical profiles of total dissolved iron, manganese, zinc, and cobalt concentrations at each station. Error bars in D represent the standard deviation of the mean (*n =* 3).

### Phytoplankton respond to cobalamin addition

Following six days of incubation, chlorophyll *a* (Chl*a*) concentrations were significantly higher in the cobalamin-only (+B_12_) treatment compared to all treatments without cobalamin (*P* < 0.02, Fig. 1d). With the addition of both iron and cobalamin (+FeB_12_), Chl*a* concentrations were 24% higher than with cobalamin alone (P = 0.01, Fig. 1d). Photosynthetic efficiency of photosystem II (F_v_/F_m_) was significantly reduced by DFB addition (P < 0.01), but otherwise, F_v_/F_m_ was not different among the other treatments (Fig. S1a). Unlike previous experiments in the Southern Ocean^8,13,14,28^, these results demonstrate that cobalamin was the primary limiting nutrient followed by iron within the phytoplankton community.

Concurrent with increases in Chl*a* concentrations, cobalamin addition resulted in greater uptake of nitrate and phosphate but not silicic acid (Fig. 1e and Fig. S1c,d). Compared to other stations in the Amundsen Sea, radiolabeled cobalamin uptake at the near surface was highest in the ASP at Station 4, indicating high cobalamin demand (Fig. 1f). At all stations, cobalamin uptake at the surface was largely driven by eukaryotes, with the ≥ 3.0 µm community accounting for 56-95% of cobalamin uptake (Fig. S2a). Within the incubations, cobalamin addition also resulted in cobalamin uptake rates that were three-to six-fold greater than in those without added cobalamin (Fig. 1h).

Cobalt uptake rates were similarly higher in the ASP compared to Station 11 and were driven by the ≥ 3.0 µm community (Fig. 1g and Fig. S2b). As cobalamin contains cobalt, its addition in the experiment resulted in a 200 pmol L^-1^ increase in dissolved cobalt concentrations (Fig. 1i). Consistent with the high cobalamin uptake rates, dissolved cobalt was consumed by 44 pmol L^-1^ (19%) in the +B_12_ treatment and 106 pmol L^-1^ (44%) in the +FeB_12_ treatment, demonstrating a cobalamin uptake capacity that far exceeds observed cobalamin concentrations (Fig. 1i).

### Recent trace metal inputs alleviate iron limitation

The initial seawater was characteristic of Antarctic Surface Water (AASW), one of the three regional endmember water types, which forms as a relatively thin and fresh surface layer from solar irradiation and seasonal ice melt, with Winter Water (WW) and modified Circumpolar Deep Water (mCDW) at deeper depths (Fig. S3)^29^. In the Amundsen Sea, mCDW flows along the continental shelf and enters ice cavities beneath the ice shelf, melting it from below^30^. Ice melt reduces the salinity of the water, which adds buoyancy and creates a “meltwater pump” that delivers trace metal-rich seawater to the surface outflow of the ice shelf^15,31^.

Near-surface dissolved iron concentrations at Station 4 were 0.01 nmol L^-1^, which was the lowest concentration measured throughout the water column among the stations (Fig. 1j). In comparison, dissolved iron concentrations within AASW in the central ASP have previously been found to be similarly low (< 0.2 nmol L^-1^) and characteristic of iron-limited regions^32–34^.

However, dissolved iron in the ASP and elsewhere is often rapidly taken up and buffered by a larger labile particulate iron pool, providing bioavailable iron to phytoplankton^34,35^. In the surface outflow of the Dotson Ice Shelf within the ASP, labile particulate iron concentrations have been observed to be 15 to 40 times those of dissolved iron^15,34^.

Compared to other stations, dissolved manganese concentrations were higher at Station 4 with a near-surface concentration of 2.64 nmol L^-1^, indicating recent high trace metal inputs from meltwater circulation (Fig. 1k). Unlike dissolved iron, manganese was higher than previous measurements in the central ASP, but less than those reported near the ice shelf outflow^33,34^.

Surface dissolved zinc concentrations were similarly higher than those at Stations 11 and 15, as were dissolved cobalt concentrations, which were higher than all other stations (Fig. 1l, m). Most dissolved cobalt was complexed to strong ligands, leaving only 7 pmol L^-1^ (19%) as labile dissolved cobalt (Fig. S2c). Copper, nickel, and cadmium concentrations at Station 4 had conservative-like profiles (Fig. S4). For zinc, copper, and nickel, these concentrations are similar to previous observations^33^.

### Diatoms and prasinophytes drive cobalamin demand

Multiple markers show that diatoms were highly prevalent, both throughout the water column at the experimental site and following six days of incubation. In the initial community, fucoxanthin (fuco), a diagnostic pigment for diatoms, was elevated compared to the pigments 19ꞌ-Hexanoyloxyfucoxanthin (19ꞌ-Hex), alloxanthin (allo), and chlorophyll *b* (chlb) (Fig. 2a-c), which are indicative of *P. antarctica*, cryptophytes, and some prasinophytes, respectively, in the region. High fucoxanthin concentrations in the initial community may also be attributed to iron-stressed *P. antarctica,* which have dominated previous summertime blooms in the ASP (Fig. 2a-c)^2,36–38^. In the initial seawater at Station 4, diatom proteins comprised 47% of the community, whereas *P. antarctica* proteins only comprised 13% (Fig. S5).

**Fig. 2.**
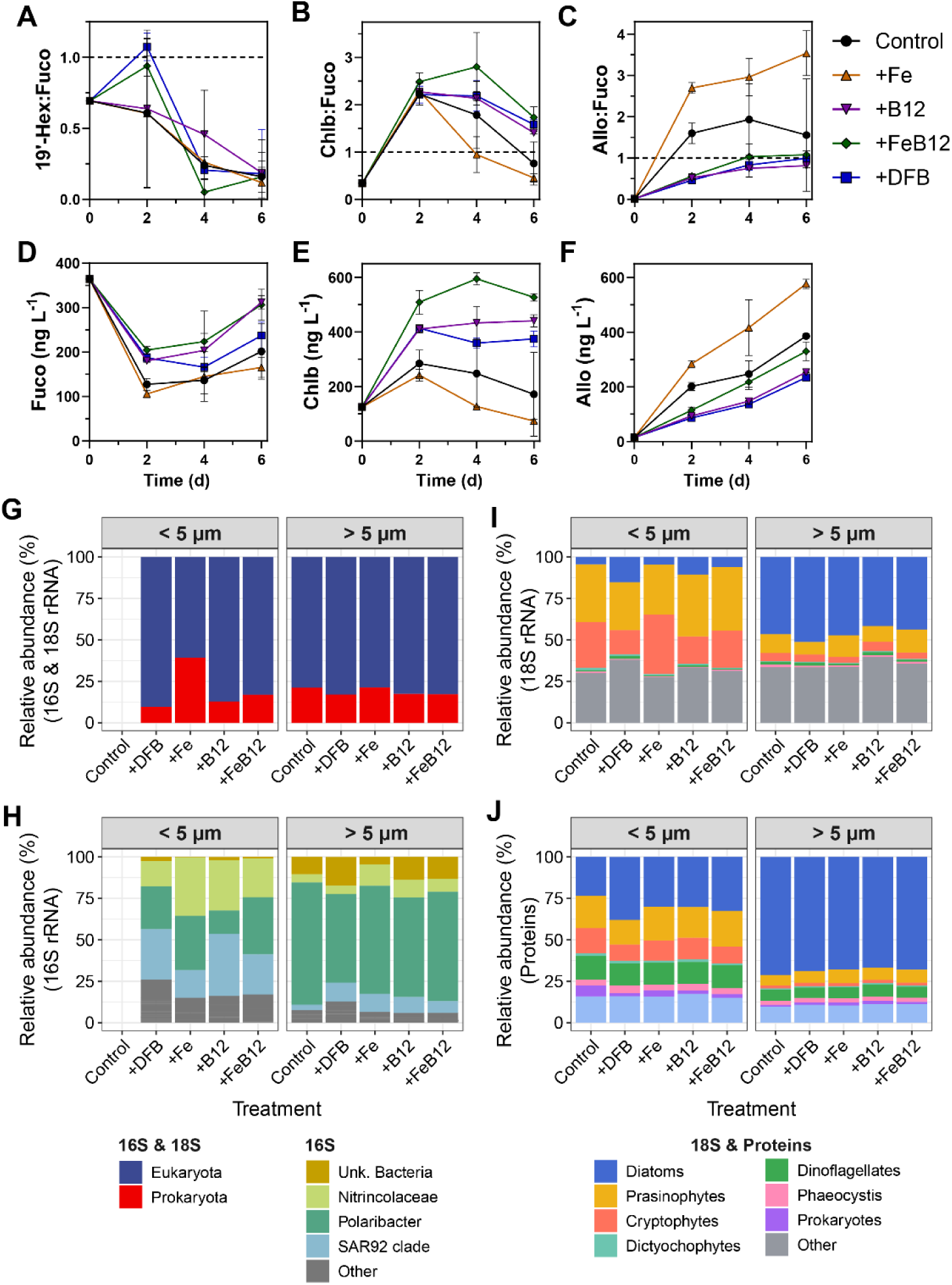
Taxonomic composition of the microbial community within the incubation experiments (A-C) Ratios of 19ꞌ-Hexanoyloxyfucoxanthin (19ꞌ-Hex), chlorophyll *b* (Chlb), and alloxanthin (Allo) to fucoxanthin (Fuco) over time. (D-F) Fuco, Chlb, and Allo concentrations (ng L^-1^) over time. 19ꞌ-Hex concentrations are shown in Fig. S1F. Relative abundances of taxa from rRNA of the (G) whole community (16S and 18S rRNA), (H) prokaryotic taxa (16S rRNA), and (I) eukaryotic taxa (18S rRNA) at the final time point. (J) Relative abundances of taxa from protein abundances. Error bars in A-F represent the standard deviation of the mean (*n =* 3).

At the final timepoint, fuco and biogenic silica concentrations, another indicator of diatom biomass^39^, were significantly higher in the +B_12_ and +FeB_12_ treatments compared to the control and +Fe treatments, demonstrating a strong response to cobalamin addition (Fig. 2d and Fig. S1b, P < 0.006). Following incubation, eukaryotes exceeded prokaryotes, representing 79-90% of rRNA and 94-99% of proteins in both size fractions, besides the +Fe treatment where 61% of rRNA was eukaryotic (Fig. 2g, j). Diatoms accounted for 42-51% of 18S rRNA within eukaryotes and 62-65% of proteins in the > 5 µm community, but only 4-15% of 18S rRNA and 23-33% of proteins in the < 5 µm community (Fig. 2i, j). From 18S rRNA, the diatom community comprised diverse genera, which were prevalent at deeper depths at Station 4 as well (Figs. S7-S9). In comparison, 19ꞌ-Hex significantly declined over time in all treatments (P < 0.0001), possibly due to preferential grazing of solitary *P. antarctica*, with *P. antarctica* representing less than 5% of rRNA and proteins at the final time point (Fig. 2A, I, J).

In the < 5 µm community, prasinophytes were also prominent. Initially, chlb concentrations were one-third those of fuco but increased in all treatments after two days and remained elevated in the cobalamin and +DFB treatments by the final time point (Fig. 2b). Like fuco, chl*b*, lutein, and zeaxanthin were significantly higher in the +B_12_ and +FeB_12_ treatments at the final timepoint, demonstrating a prasinophyte response to cobalamin addition (P < 0.0001) (Figs. 2e and Fig. S6). In the < 5 µm community, prasinophytes were 29-38% of the 18S rRNA and 17-24% of proteins with > 98% of rRNA reads attributed to *Pyramimonas*, which are mixotrophic (Fig. 2i, j)^40^.

Rather than responding to cobalamin addition, significant increases in alloxanthin show that cryptophytes increased solely in the +Fe treatment (P < 0.001) (Fig. 2f). Increases were not observed in the +FeB_12_ treatment, indicating that they were outcompeted when both nutrients were replete. In the < 5 µm community, cryptophytes were prevalent as well, accounting for 15-36% of the 18S rRNA and 11-15% of proteins (Fig. 2i, j). Consistent with increases in alloxanthin, the highest rRNA relative abundances were in the control and +Fe treatments. More than 98% of cryptophyte-assigned rRNA was attributed to *Geminigera cryophila*.

The unique response in *G. cryophila* indicates that it has different iron and cobalamin requirements or acquisition strategies. Previous cobalamin quota measurements in the cryptophyte *Rhodomonas salina* shows that its cobalamin requirement is low^41^, and as mixotrophs, cryptophytes may satisfy their cobalamin requirement through bacterivory, leveraging their motility to improve encounter rates^42^. However, bacterivory may not provide sufficient iron, leading to iron limitation instead. Notably, cryptophyte populations in the Southern Ocean have been rising from 1997-2023, indicating more favorable growth conditions and an increased ability to compete for resources under modern conditions^3^.

### Prokaryotic capacity for cobalamin synthesis is limited

Prokaryotic cell abundances were 4.05 ± 0.38 x 10^5^ cells mL^-1^ in the initial seawater and only showed reductions in the +DFB treatment compared to the control treatment (2.37 ± 0.14 x 10^5^ cells mL^-1^, P = 0.18, Fig. S10). With Archaea representing less than 0.1% of both rRNA and proteins, 10-21% of rRNA was bacterial, except the +Fe treatment in the < 5 µm community where bacteria were 39% of rRNA, indicating stimulation of bacterial activity with iron as observed in other systems (Fig. 2g)^43,44^. Bacterial proteins comprised only 4.2-6.2% of the < 5 µm community and 1.2-1.5% of the > 5 µm community (Fig. 2j), with the overall community dominated by eukaryotic phytoplankton as previously described. At the experimental site, bacterial proteins at 40 m were only 54% of the 0.2-3 µm community and < 7% in larger fractions (Fig. S11).

The bacterial community in the > 5 µm community was dominated by the genus *Polaribacter* (53-74%) with seven additional unidentifiable taxa (ASVs) accounting for 4-17% (Fig. 2h). *Polaribacter* were also present in the < 5 µm community (14-35%), but taxa assigned to the family Nitrincolaceae (formerly Oceanospirillaceae)^45^ and the SAR92 clade were also abundant, representing 15-37% of this size fraction. In the < 5 µm community, high abundances of *Polaribacter* and Nitrincolaceae correspond to the increased relative abundances of prokaryotes overall in the +Fe treatment (Fig. 1g). All of these taxa similarly dominated the bacterial community during previous *Phaeocystis* blooms in the ASP^37,46,47^.

Nitrincolaceae have previously been identified as cobalamin producers in the region^13,46^. However, with currently available genomes, only one-third of Nitrincolaceae and *Polaribacter* are possible cobalamin producers (Fig. S12a, b)^48^. No SAR92 clade bacteria appeared to contain a majority of the pathway. The cobU gene enables currently identified mechanisms of cobalamin scavenging in model bacteria as an alternative source but was largely absent as well^49^.

A key use of cobalamin is for the production of methionine by methionine synthase (MetH), although some organisms possess a cobalamin-independent but less efficient form (MetE)^50^. While the majority of examined genomes have either *metH* or both genes, 18% of *Nitrincolaceae* possess neither gene, suggesting a reliance on exogenous sources of methionine despite cobalamin synthesis capabilities (Fig. S12c)^51^.

Altogether, prokaryotic abundances relative to eukaryotes were low, likely due to a lack of dissolved organic carbon in early summer^52^. Moreover, genomic capacity for cobalamin synthesis within the prokaryotic community was also limited. With competition for cobalt, a lack of cobalamin-producing prokaryotes limits cobalamin supply, creating strong competition for cobalamin among both non-producing bacteria and phytoplankton (Fig. S2).

### Proteomic responses align with altered nutrient states

Within diatoms, molecular indicators of nutrient stress established by culture studies displayed expected differences among the different treatments in both size fractions (Fig. 3a, b). In the +B_12_ and +FeB_12_ treatments, indicators of cobalamin stress were low compared to all treatments. These include cobalamin acquisition protein 1 (CBA1), a shift from MetE to MetH, methionine adenosyltransferase (MetK), and serine hydroxymethyltransferase (SHMT)^53^. Abundances of these proteins were often highest in the +Fe treatment.

**Fig. 3.**
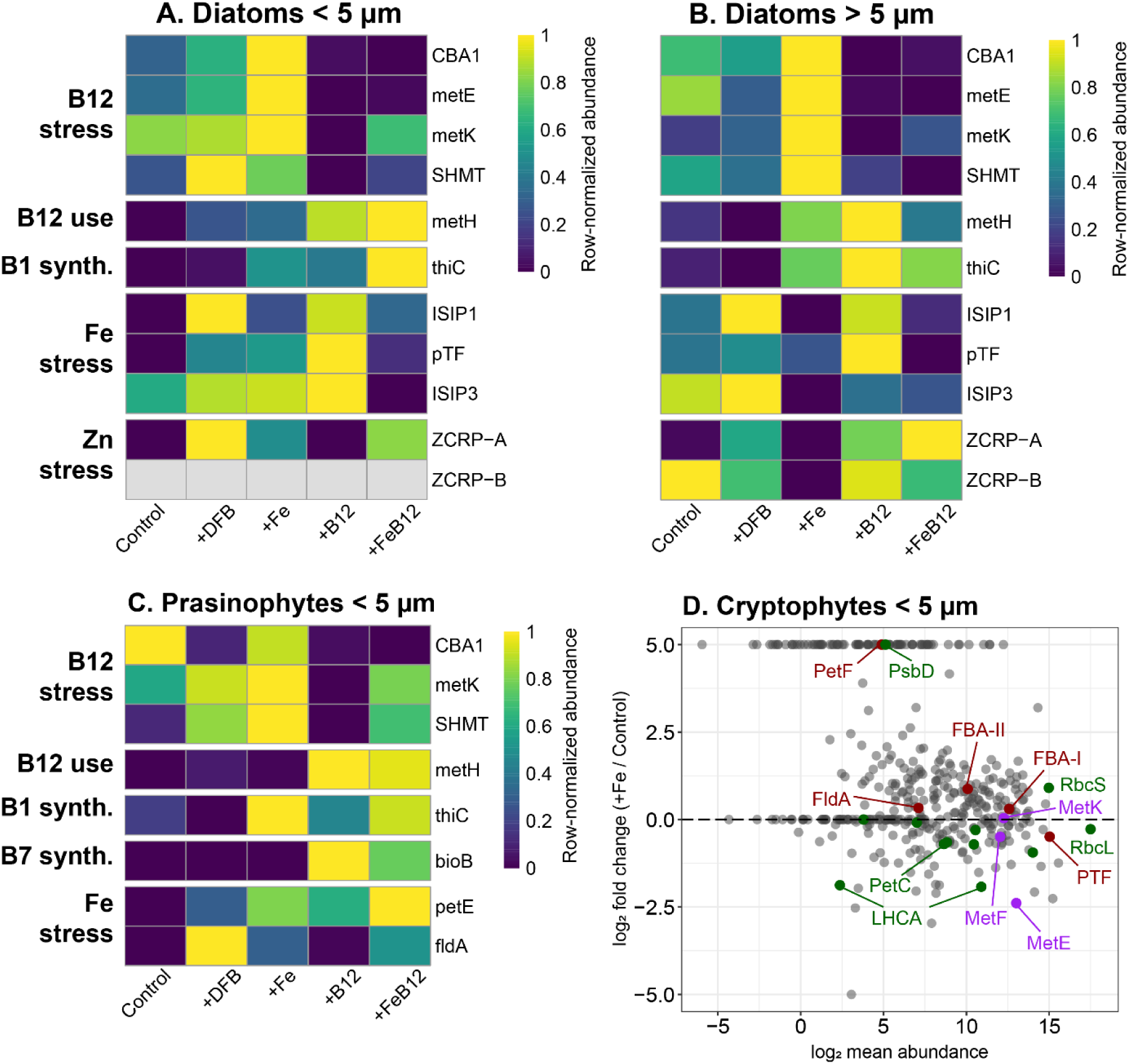
Proteomic responses among treatments within the (A) < 5 µm diatom community, (B) > 5 µm diatom community, and (C) < 5 µm prasinophyte community at the final time point. Proteins are categorized based on nutrient stress or function, and abundances are row-normalized. (D) Mean-difference (MA) plot for the cryptophytes in the < 5µm community comparing the +Fe and control treatments. Proteins are colored as related to cobalamin-related (purple), iron-responsive (red), or photosynthesis-related (green).

Iron addition, with or without cobalamin, largely resulted in lower abundances of iron stress induced proteins (ISIP1, ISIP2A/PTF, and ISIP3; Fig. 3a, b)^54^. Concordantly, cobalamin alone and DFB addition resulted in greater abundances of these iron stress proteins, indicating that cobalamin addition drove the community towards iron stress. In larger diatoms with greater Zn requirements^55^, abundances of zinc-responsive proteins (ZCRP-A, ZCRP-B) increased with cobalamin and DFB addition and were lower with iron addition, indicating zinc stress occurred when cobalamin limitation was alleviated or when Zn was chelated by DFB^56^.

In the < 5 µm community where prasinophytes were prevalent, proteomic responses were similar to those in diatoms (Fig. 3c). Cobalamin addition resulted in low CBA1 abundances. In the +B_12_ treatment, MetK and SHMT abundances were also low, but elevated in the +FeB_12_ treatment. As few prasinophytes possess *metE*, no prasinophyte MetE proteins were detected, but MetH was highest with cobalamin addition^57,58^. Both flavodoxin (FldA) and plastocyanin (PetE) are associated with iron stress^54^, and while FldA was highest in the DFB treatment, PetE abundances were highest in the treatments containing iron and/or cobalamin. Continued use of PetE rather than cytochrome c_6_, which was undetected, is similar to observations in the iron-limited Northeast Pacific and may be an adaptation to chronically low iron conditions^59^.

In both diatoms and prasinophytes, increased abundances for the vitamin B_1_ (thiamine) synthesis protein, phosphomethylpyrimidine synthase (ThiC), were observed with iron and/or cobalamin addition (Fig. 3)^60^. Prasinophytes also displayed increased abundances of the vitamin B_7_ synthesis protein, biotin synthase (BioB), with cobalamin addition^61^. The increased expression of both proteins indicates an increased demand for other B-vitamins after iron or cobalamin addition.

To examine the increase in cryptophytes in the +Fe treatment (Fig. 2f, i), differential protein abundances from cryptophytes in the < 5 µm community between the +Fe and control treatments were evaluated (Fig. 4d). Corresponding to reduced iron stress, shifts from FldA to ferredoxin (PetF) and from class I to class II fructose bisphosphate aldolase (FBA) were observed with iron addition^54^. Phytotransferrin (PTF/ISIP2A) abundances were also lower. MetH was not detected; however, MetE abundances declined with iron addition along with MetF, which provides 5-methyltetrahydroflate to MetH and MetE^62^. MetK was unchanged. Most other proteins involved in photosynthesis had reduced abundances, particularly light-harvesting complex chlorophyll-binding proteins (LHCA) (Fig. 3d). The large subunit of RuBisCo (RbcL) and components of the photosynthetic electron transport chain had reduced abundances as well. However, the small subunit of RuBisCo (rbcS), iron-responsive PetF, and a component of photosystem II (PsbD) showed increased abundances with iron.

**Fig. 4.**
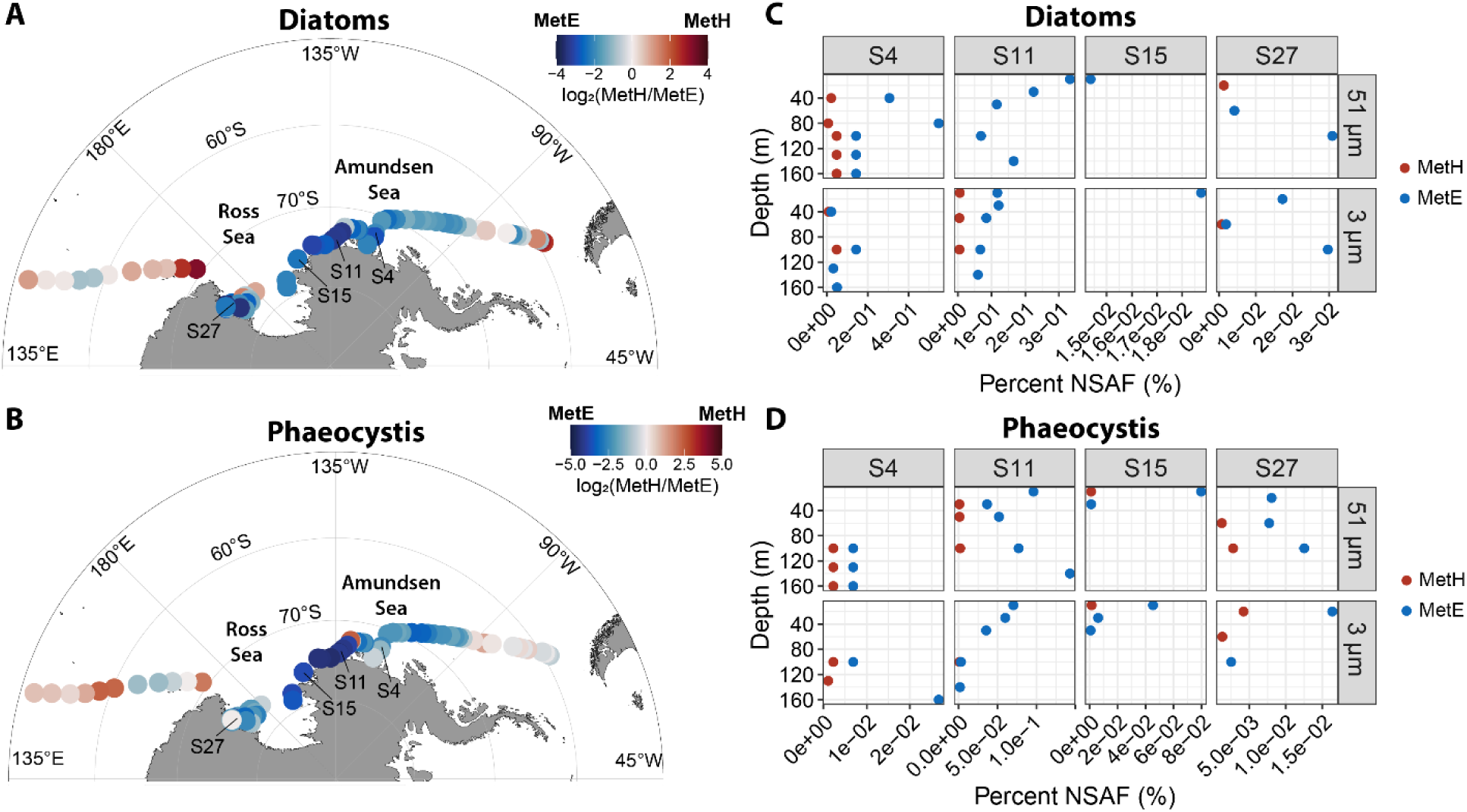
MetH versus MetE protein abundances in diatoms and *Phaeocystis.* (A, B) The log_2_ ratio of MetH versus MetE from surface underway samples. (C, D) Vertical profiles of MetH (red) or MetE (blue) expressed as percent normalized spectral abundance factor (NSAF) within each taxonomic group for the > 51 µm and 3-51 µm communities.

Although cryptophytes possess the *metH* gene^58^, the lack of MetH use likely leads to a dramatically lower cobalamin requirement and underlies their ability to grow in the +Fe treatment. CBA1 was also not detected, indicating that cryptophytes may leverage a different cobalamin acquisition strategy, such as bacterivory, to meet their low cobalamin requirement. The observed shifts in iron-responsive genes and down-regulation of photosynthesis are also consistent with impairment of their photosynthetic apparati due to iron limitation.

### Widespread cobalamin stress in the Southern Ocean

During the same expedition, near-surface seawater was collected for metaproteomics while transiting from Chile to the Amundsen and Ross Seas and then to Australia, providing a nearly 10,000 km long transect (*n =* 80, Data S1). Consistent with previous observations of the seasonal sea ice zone^3^, diatoms in surface waters were more abundant than *Phaeocystis*, except 10 samples from the western Amundsen/Eastern Ross Seas (Fig. S5). *Phaeocystis* protein abundances were also elevated throughout the upper water column at stations in that region (Fig. S11).

Within these samples, the MetH:MetE ratio in diatoms and *Phaeocystis* was used to evaluate cobalamin stress (Fig. 4A,B). In both taxa, MetE exceeded MetH abundances in 79% of samples. Protein abundances throughout the upper water column (< 160 m) at four coastal stations also showed greater MetE abundance compared to MetH (Fig. 4c, d). In diatoms, the strongest shift towards MetE was found in the coastal regions of the Amundsen and Ross Seas. Diatom CBA1 abundances exhibited a similar pattern with highest abundances in the Ross Sea (Fig. S13).

Overall, 49% of samples had CBA1 abundances that exceeded those at Station 4, where cobalamin limitation was detected. In the western offshore regions, MetE exceeded MetH, but MetH was more abundant in eastern offshore samples north of 70°S, either from geographical or temporal differences (Fig. 4a, b).

*P. antarctica* is an important component of Antarctic coastal ecosystems, yet the extent to which natural *P. antarctica* populations experience primary cobalamin limitation has remained an open question. Their abundances within the experiment were low, potentially due to grazing, stratification, and recent trace metal inputs that favor diatoms^63^. However, *P. antarctica* often blooms early within Antarctic polynyas upon sea ice retreat and is followed by diatom blooms, which may have occurred here as the polynya studied here developed at least one month prior to the experiment (Fig. S14)^64^.

Notably, *P. antarctica* did not increase in biomass upon cobalamin addition here or in a prior experiment in the Ross Sea (Fig. 2i, j)^14^. Yet, laboratory studies with *P. antarctica* have demonstrated cobalamin’s benefit for growth and MetE use to reduce demand under low cobalamin conditions^57^. Indeed, cobalamin stress in *P. antarctica* was evident with MetH:MetE abundances similar to those in diatoms, but *Phaeocystis* MetE did not as strongly exceed MetH expression in the eastern Ross Sea, implying that *P. antarctica* is less susceptible to cobalamin limitation (Fig. 4a, b).

Besidess MetE use, *Phaeocystis* forms symbiotic relationships with bacteria in their colonial matrix that allow them to obtain B-vitamins^65^. Increased iron inputs also cause *P. antarctica* to transition from its solitary to colonial morphotype supporting the formation of these symbiotic relationships or bacterivory early in the season during polynya formation^57,66,67^. Together, these adaptations may reduce cobalamin stress in *P. antarctica* and allow them to obtain cobalamin more readily than diatoms when cobalamin and bacterial abundances are lower^52^.

### Conclusions

Warming is increasing iron supplies through increased ice melt in coastal regions of the Southern Ocean as well as reducing demand in phytoplankton through increased iron use efficiencies^15–17,31,68–70^. With decreased iron stress, demand for cobalamin increases; however, cobalamin availability may not be rising accordingly, with low bioavailable cobalt that is largely consumed by eukaryotes and a microbial community that is largely not capable of cobalamin synthesis^21^. As a result, the increases in Chl*a* and macronutrient drawdown with cobalamin addition observed here demonstrate primary cobalamin limitation rather than iron limitation or iron-cobalamin colimitation as previously shown^8,13,14,28^.

Although cobalamin limitation was evident in diatoms and prasinophytes, cryptophytes were iron limited. Differences in cobalamin stress were also apparent between diatoms and *Phaeocystis* as well, underscoring differences in cobalamin requirements and acquisition strategies among different phytoplankton taxa. Thus, consideration of cobalamin as a primary, secondary, or colimiting nutrient as well as different taxon-specific responses is necessary to project changes to phytoplankton community structure and productivity in Antarctic polynyas, which have a significant role in Earth’s carbon cycle (Table S1)^6,7^.

## Methods

### Overview of sample collection

Sample collection occurred during the CICLOPS (Cobalamin and Iron Co-limitation of Phytoplankton Species) expedition (NBP18-01), which took place onboard the RVIB *Nathaniel B. Palmer* from 11 December 2017 – 3 March 2018 and transited from Chile to the Amundsen and Ross Seas and Tasmania, Australia (Fig. 4a, b). Throughout the cruise, samples were collected for metaproteomics from the ship’s underway seawater system. Water column samples were also collected at Station 4, 10, 11, and 15 within the Amundsen Sea (Fig. 1a-c). At these stations, seawater was collected using a trace metal clean rosette equipped with 8 L Niskin-X bottles and analyzed for dissolved trace metal concentrations, cobalamin and cobalt uptake rates. At these stations as well as another station in the Ross Sea (Station 22), samples for RNA and metaproteomics were collected with McLane pumps.

### Experimental setup

At Station 4 (72° 45.00 S, 116° 00.03 W, Fig. 1a0c) on December 31^st^, 2017, a trace metal clean incubation experiment was performed. Seawater was collected with a towfish and pumped directly into a trace metal clean van with a Teflon diaphragm pump into clean 50 L carboys. Seawater was then dispensed into 1 L polycarbonate bottles, 8 L polycarbonate bottles for RNA and proteins at the final time point or immediately sampled for initial (T0) measurements. All bottles were rinsed with the same seawater before filling.

Triplicate 1 L incubation bottles were either unamended (control) or amended with iron (+Fe, 1 nmol L^-1^), the chelator desferrioxamine B (+DFB, 100 nmol L^-1^), cobalamin (+B_12_, 200 pmol L^-1^), or iron and cobalamin (+FeB_12_) at the same concentrations as the individual additions. The cobalamin stock solution was treated with Chelex 100 resin to remove contaminating metals ^71^. The 8 L polycarbonate bottles for RNA and proteins at the final time point were similarly amended but with one replicate each.

At the initial time point, samples were collected for chlorophyll *a*, macronutrient, biogenic silica, dissolved Co, and flow cytometry. At days 2, 4, and 6, additional samples were collected for phytoplankton pigments, cobalamin uptake, and photosynthetic efficiency (F_v_/F_m_). At day 6 (the final time point), samples for RNA and proteins were collected from the 8 L bottles by filtering seawater onto 5 µm polycarbonate and GF/F filters in series. Three-quarters of each filter was reserved for proteomics and one-quarter of each filter was reserved for ribosomal RNA.

### Sea ice concentration and bathymetry maps

Sea ice concentration data at 12.5 km resolution from the Advanced Microwave Scanning Radiometer (ASMR) instruments were downloaded using NASA Earthdata Search. Sea ice concentration and bathymetry maps were generated with ggOceanMaps^72^.

### Chlorophyll a

Seawater (100 mL from incubation bottles, 280 mL for T0) was filtered onto GF/F filters under gentle vacuum pressure and frozen at –20 °C until analysis. Chlorophyll *a* extraction was performed using 90% acetone at –20 °C for 24 hours and measured via in vitro fluorometry on a Turner Designs 10-AU fluorometer using the acidification method^73^.

### Photosynthetic efficiency (F_v_/F_m_)

The maximum quantum yield of photosystem II (F_v_/F_m_) was measured with a Phyto PAM analyzer (Walz). Samples were dark acclimated for 30 min at 0 °C to obtain minimal fluorescence, then subjected to a short pulse of saturating light (>3000 µmol photons m^-2^ s^-1^ for 0.8 ms) to obtain maximum fluorescence. Measurements were blank corrected with 0.2 µm filtered seawater.

### Macronutrient concentrations

Seawater was filtered with 0.2 µm polyethersulfone filters into acid-washed 60 mL high-density polyethylene bottles and frozen until analysis. Samples were analyzed with a Technicon Autoanalyzer II by Joe Jennings at Oregon State University following the methods described in Gordon, et al. ^74^. Technicon AutoAnalyzer II components were used to measure phosphate, and Alpkem rapid flow analyzer (RFA) 300 components were used to measure nitrate+nitrite and silicic acid.

### Biogenic silica

Seawater (250 mL from incubation bottles, 1060 mL for T0) was filtered onto 0.6 µm polycarbonate filters under gentle vacuum pressure. Filters were placed in glassine envelopes and dried at 65 °C for approximately 24 h. For analysis, filters were extracted in 0.2N NaOH for 45 minutes at 95 °C. Samples were placed on ice and neutralized with 1N HCl to stop the extraction. Concentrations of silicic acid were determined using the colorimetric method described in Strickland and Parsons ^75^. Precautions were made to avoid contamination by using plasticware and Milli-Q water.

### Phytoplankton pigments

Phytoplankton pigments were measured with high performance liquid chromatography (HPLC) with a modification of the Zapata et al. ^76^ method described by DiTullio and Geesey ^77^. Briefly, seawater was gently filtered under low vacuum pressure onto GF/F filters and frozen at –80 °C until analysis onshore. Samples were extracted in acetone and analyzed with an Agilent 1100 HPLC system equipped with an autosampler, photodiode array, and fluorescence detectors.

### Dissolved trace metals

Total dissolved Fe, Mn, Zn, Cu, Cd, and Ni concentrations were collected and analyzed as described by Kell et al. ^78^. Briefly, seawater was collected at various depths with 8 L Niskin-X bottles on a trace metal clean rosette suspended from a Kevlar line and pressure-filtered with high-purity N_2_ gas through acid-washed 0.2 µm Supor™ polyethylsulfone filters. Samples were acidified to approximately pH 1.7 with HCl (Optima™) within seven months of sampling and stored acidified for over one year prior to analysis.

Quantification was performed using the isotope dilution method where 15 mL of seawater was enriched with stable isotopes. Seawater was then concentrated to 500 µL with a seaFAST pico containing a NOBIAS-chelate PA1 resin column. Multi-elemental analysis was then performed on a Thermo Scientific iCAP-Q.

Dissolved cobalt was collected and measured described by Chmiel et al. ^21^ using the method established by Saito and Moffett ^79^ and modified by Saito et al. ^80^ and Hawco et al. ^81^. Briefly, filtered seawater was collected in 60 mL low-density polyethylene (LDPE) bottles and stored at 4°C until analysis via cathodic stripping voltammetry (CSV) at sea within three weeks. CSV analysis was conducted using Metrohm 663 VA and *μ* Autolab III systems equipped with a hanging mercury drop working electrode. For total dissolved cobalt, samples were first UV-irradiated in quartz tubes for 1 h in a Metrohm 705 UV Digester to destroy natural ligand-bound cobalt complexes, whereas labile dissolved cobalt samples were not UV-irradiated.

### Cobalt and cobalamin uptake

Trace metal clean seawater was dispensed into clean 60 mL polycarbonate bottles and amended with 0.1 pmol L^-1^ of either ^57^CoCl_2_ or ^57^Co-labeled cobalamin. Bottles were placed in an on-deck incubator with flow-through seawater and screening to achieve approximately 20% surface irradiance for 24 hours. Following incubation, seawater was filtered onto 0.2 µm and 3.0 µm filters separately. Filters were then stored in petri dishes until γ-ray counting. Without cobalamin concentrations, only percent cobalamin uptake can be calculated; estimated cobalamin uptake rates for experimental treatments assume that starting background cobalamin concentrations were 1 pmol L^-1^ in the treatments without cobalamin and 200.1 pmol L^-1^ for treatments with cobalamin. Later time points for treatments with cobalamin estimate drawdown of cobalamin based on the decline in dissolved cobalt concentrations.

### Prokaryotic cell abundances

Prokaryotic cell abundances were measured with flow cytometry. Samples (1 mL) were fixed for 10 min with a mix of both paraformaldehyde and glutaraldehyde (1 % and 0.05 % final concentration, respectively), frozen in liquid nitrogen, and stored at −80°C until analysis. After thawing for analysis, samples were stained with SYBR Green I using a 1000x dilution of the stock solution for 15 min at room temperature. Cell concentrations were assessed using a BD FACSVerse flow cytometer equipped with a 488 nm Ar laser and standard set of optical filters and analyzed with FCS Express 6 Flow v 6.06.0025 (DeNovo Software). Prokaryotes were discriminated against other particles based on scatter and green fluorescence from SYBR Green I.

### Metatranscriptomics

Samples for metatranscriptomics were collected during the cruise with McLane pumps within the Amundsen and Ross Seas. With depths ranging from 10 m to 464 m, seawater was filtered in series onto 51 µm PETEX (Sefar), 3 µm Versapor® acrylic, and 0.2 µm Supor® PES filters, and ¼ of the filter was frozen at −80°C for RNA. Once onshore, RNA was extracted from the filters on an Eppendorf epMotion 5075 TMX with the Machery-Nagel NucleoMag RNA kit.

From the 0.2 µm filters, rRNA was removed using siTOOLs riboPOOLs with a 6:1:1 mixture of the pan-prokaryote, pan-plant, and pan-mammal riboPOOLs following the manufacturer’s instructions for low input. rRNA depletion was confirmed with all samples on an Agilent TapeStation 2200 with High Sensitivity RNA ScreenTape. cDNA was then synthesized using the Ovation RNA-Seq System V2 kit (NuGEN) using both poly-A and random hexamer primers followed by fragmentation with a Covaris® E210 focused-ultrasonicator targeting 300 bp fragments. cDNA was then purified with Agencourt RNAClean XP beads, and library preparation was performed with the Ovation Ultralow System V2 with unique dual indexes (NuGEN). After end repair, ligation, and amplification, the libraries were assessed on an Agilent TapeStation 2200 with High Sensitivity DNA ScreenTape.

From the 3 and 51 µm, samples with total RNA greater than 1 µg, 800 ng of total RNA was used for preparing poly-A libraries with an Illumina Stranded mRNA Prep Ligation kit (Illumina), following the manufacturer’s protocol. For the 3 and 51 µm samples with total RNA less than 1 µg, 20 ng of total RNA was used as input for the SMART-Seq v4 Ultra Low Input RNA kit (Takara Bio USA. Inc). The resulting double-stranded cDNA was then fragmented using a Covaris E210 system with the target size of 300 bp. Libraries were prepared from fragmented double-stranded cDNA using an Illumina Stranded mRNA Prep Ligation kit (Illumina). Resulting libraries from all samples were sequenced on an Illumina NovaSeq 6000 with a S4 flow cell (PE150).

Raw reads were trimmed for quality and removal of Illumina adaptors. Ribosomal RNA sequences were also removed with Ribopicker v0.4.3^82^. Trimmed and filtered reads were then used for assembly into contigs, and abundances were quantified by mapping these reads to the assembly. Both assembly and read mapping were performed with CLC Bio Genomics Server v21.0.3. Gene prediction was performed with FragGeneScan v1.16, and rRNA removal was performed again with Ribopicker^83^. Predicted proteins were further filtered to remove those less than 10 amino acids long or with greater than or equal to 20% stop codons.

With the final assemblies, taxonomic annotation of each protein coding gene was assigned via DIAMOND BLASTP alignments using PhyloDB v1.076 as a reference database^13^. Final taxonomic assignments were based on highest Lineage Probability Index values^84^. For functional annotation, DIAMOND BLASTP alignments were performed with the Kyoto Encyclopedia of Genes and Genomes (KEGG; Release 94.1)^85^. KEGG Ortholog (KO) assignment was performed with KofamKOALA, which uses hmmsearch against KOfam, an HMM database of KOs^86^. E-value cutoffs of 1e-5 were used throughout.

### Metaproteomics (experimental and underway samples)

For experimental incubation samples, approximately 7 L of seawater was filtered in series onto 5 µm polycarbonate and GF/F filters. One-fourth of each filter was separated, placed into cryovials and frozen for rRNA sequencing with the remaining piece reserved for proteomics. The remaining slices of the 5 µm filters were also put into cryovials, and the remaining slices GF/F filters were folded and wrapped in Al foil. Samples were also collected from the ship’s underway seawater supply directly onto 142 mm GF/F filters (Data S1). Filters were frozen at −80°C until extraction.

Proteins were extracted from each filter with 8 mL of lysis buffer (50 mM HEPES pH 8.5, 1% SDS). All reagents were made with HPLC-grade (Optima™) water. Filters were incubated with the lysis buffer at 95°C for 12 min and then room temperature for 30 min while shaking. Filters were then removed from tubes, and the remaining solution was syringe filtered with a 5 µm filter. Syringes were rinsed with an additional 900 µL of lysis buffer to collect remaining material. Following, samples were centrifuged at 3,220 x g for 30 min. The supernatant was then concentrated to 350 to 650 µL with Vivaspin 6 centrifugal concentrators (5,000 MWCO) at 3,220 x g then quantified with the Pierce Micro BCA™ Protein Assay Kit.

35-50 µg of protein was subsampled and diluted with lysis buffer to a volume of 200 µL. 2 µL of Benzonase Nuclease (> 90% purity, Millipore) was then added to each sample for 30 min at 37°C. Samples were then reduced with 10 µL of 100 mM dithiothreitol (DTT) for 30 min at 45°C, cooled to room temperature, then alkylated with 20 µL of 400 mM iodoacetamide (IAA) in the dark for 30 min at room temperature. Another 20 uL of 100 mM DTT was then added and incubated for 10 min at room temperature to quench residual IAA.

Sample cleanup was performed with Sera-Mag Carboxylate-Modified Magnetic Beads and SpeedBeads (Cytiva) according to the manufacturer’s instructions based on Hughes, et al. ^87^ and Hughes, et al. ^88^. Briefly, proteins were bound to 350-500 µg of beads at a final concentration of 50% ethanol, then washed twice with 80% ethanol. Proteins were digested overnight with 1.6 µg of mass spectrometry grade trypsin (Promega) at 37°C. Beads were then replaced with new beads, and binding was initiated with a final concentration of 95% acetonitrile. Peptides were then washed again with 100% acetonitrile (MeCN) and eluted in 2% DMSO. Peptides were further purified with Pierce™ C18 spin columns according to the manufacturer’s protocol, then dried completely with a SpeedVac (Thermo Scientific), and resuspended in Buffer B (0.1% formic acid, 2% MeCN).

Approximately 2 µg of tryptic peptides were analyzed with liquid chromatography coupled with tandem mass spectrometry (LC/MS/MS) where a Thermo Vanquish Neo HPLC reverse phase chromatography system was paired with a Thermo Scientific Orbitrap-Astral mass spectrometer using a Thermo Flex nanospray source. The peptides were eluted on a reverse phase C18 column (0.1 x 500 mm ID, 2 µm particle size, 120 Å pore size, C18 Reprosil-Gold, Dr. Maisch GmbH) using a 500 nL min^-1^ flow rate. All solvents were prepared with Fisher Optima grade reagents.

Liquid chromatography was a nonlinear 120-minute gradient from 5%-95% buffer B, where buffer A was 0.1% formic acid. Data-independent acquisition (DIA) mode was used with a precursor range of 380 m/z to 980 m/z on the Orbitrap at 480K resolution. MS2 scans were performed with 3 m/z MS2 non-overlapping windows on the Astral analyzer with an injection time of 3.5 ms and a range of 150-2000 m/z.

Peptide and protein identification was performed using a three-stage workflow within DIA-NN v.2.3.2^89^. In the first stage, a calibration library was generated from a subset of raw files searched against the metatranscriptome assembly. This library was then supplied as a reference for the second stage, in which InfinDIA mode was used to pre-search all the raw files and build an empirical spectral library from confidently identified precursors. In the third and final stage, all raw files were quantitatively searched against this spectral library using standard DIA-NN scoring and quantification to generate the final protein group, gene, and precursor identification and quantification matrices.

Searches were performed at a 1% FDR (precursor and protein q-value ≤ 0.01), with tryptic specificity (cleavage after K/R), up to one missed cleavage, peptide lengths of 7–35 residues, precursor m/z 380–980, charge states 2–4, and fragment m/z 150–2000. Oxidation of methionine was included as a variable modification, with N-terminal methionine excision enabled, and mass accuracy tolerances were calibrated automatically at each stage (MS1/MS2 recalibration with an initial 25 ppm window). RT-dependent cross-run alignment and match-between-runs–style reanalysis were enabled at each of the three stages to improve consistency of identification and quantification across the run set. Protein inference was enabled only for the final quantitative search. edgeR v4.0.16 was then used to normalize abundances within taxonomic groups among the experimental treatments^90^.

### Metaproteomics (vertical profiles)

Along with RNA described above, samples were collected during the cruise with McLane pumps. Samples from stations 4, 11, 15, and 27 at ≤ 160 m were used here. Seawater was filtered in series onto 51 µm PETEX (Sefar), 3 µm Versapor® acrylic, and 0.2 µm Supor® PES filters, and one-half of the filter was frozen at −80°C for proteomics. As described for the other metaproteomic samples, samples were lysed, concentrated, and treated with benzonase nuclease.

Extracted proteins were then purified from SDS detergent, reduced, alkylated, and digested with trypsin while embedded within a polyacrylamide tube gel^91^. All plastic microcentrifuge tubes were ethanol-rinsed and dried prior to use. A gel premix was made by combining 1 M Tris HCl (pH 7.5) and 40% Bis-acrylamide L 29:1 (Acros Organics) at a ratio of 1:3. The premix (103 µL) was combined with 50–100 µg of the extracted protein sample, Tris-EDTA, 7 µL 1 % APS, and 3 µL of TEMED (Acros Organics) to a final volume of 200 µL. After 1 h of polymerization at room temperature, 200 µL of gel fix solution (50% ETOH, 10% acetic acid) was added to the top of the gel and incubated at room temperature for 20 min. Liquid was then removed and the tube gel was transferred into a new 1.5 mL microcentrifuge tube containing 1.2 mL of gel fix solution before incubating at room temperature at 350 rpm for 1 h. Gel fix solution was removed and replaced with 1.2 mL of destain solution (50% MeOH, 10% acetic acid) and incubated at 350 rpm and room temperature for 2 h. Liquid was removed, and gels were cut up into 1 mm cubes and added back to tubes containing 1 mL of 50:50 acetonitrile:25 mM ammonium bicarbonate (ambic) and incubated for 1 h at 350 rpm and room temperature. Liquid was removed and replaced with fresh 50:50 acetonitrile:ambic solution and incubated at 16°C and 350 rpm overnight. The above step was repeated for 1 h the following morning. Gel pieces were then dehydrated twice in 800 µL of acetonitrile for 10 min at room temperature and dried for 10 min in a SpeedVac.

Proteins were reduced in 600 µL of 10 mM DTT, 25 mM ambic at 56°C and 350 rpm for 1 h. The volume of unabsorbed DTT solution was measured prior to removal. Gel pieces were washed with 25 mM ammonium bicarbonate, and 600 µL of 55 mM IAA was added to alkylate proteins at RT and 350 rpm for 1 h. Gel cubes were then washed with 1 mL ambic for 20 min and 350 rpm at RT. Acetonitrile (1 mL) dehydrations and SpeedVac drying were repeated as described above. Trypsin was added in an appropriate volume of 25 mM ambic to rehydrate and submerse gel pieces at a concentration of 1:20 µg trypsin:protein. Proteins were digested overnight at 350 rpm at 37°C. Unabsorbed solution was removed and transferred to a new tube. 50 µL of peptide extraction buffer (50 % acetonitrile, 5% formic acid in water) was added to gels, incubated for 20 min at RT, then centrifuged at 14,100 × *g* for 2 min. The supernatant was collected and combined with the corresponding unabsorbed solution. The above peptide extraction step was repeated again, combining corresponding supernatants. Combined digested peptides were centrifuged at 14,100 × *g* for 20 min, and supernatants were transferred into a new tube and dehydrated down to approximately 20 µL in the SpeedVac. Total digested peptides were quantified with the Bio-Rad DC protein assay.

Approximately 2 µg of purified peptides was then analyzed with LC/MS/MS using a Thermo Dionex UltiMate 3000 RSLCnano LC system with an additional RSLCnano pump run in online two-dimensional active modulation mode coupled to a Thermo Fusion Orbitrap mass spectrometer^92^. The mass spectrometer collected MS1 spectra at 240 K resolution between 380 and 1580 *m*/*z* with an AGC target of 4.0 × 10^5^. MS2 spectra were collected on the ion trap in data-dependent acquisition (DDA) mode with a cycle time of 2 s between master scans. The maximum injection time was 50 ms with the option to inject ions for all available parallelizable time. The CID collision energy was set at 30% and a quadrupole isolation window of 1.6 *m*/*z*. Monoisotopic peak determination was set to peptide and to include charge states from 2 to 10 with a mass tolerance of ± 3 ppm. Dynamic exclusion time was 20 s for LC–MS and 5 s for LC × LC–MS to best match the peak widths obtained for both systems.

Peptide spectra matches were performed using the SEQUEST algorithm against the metatranscriptome within Proteome Discoverer v.2.1 (Thermo Fisher Scientific) with a fragment tolerance of 0.6 Da and parent tolerance of 10 ppm. Identification criteria consisted of a peptide threshold of 95 % (false discovery rate (FDR) = 0.1%) and protein threshold of 99% (1 peptide minimum, FDR = 0.8%) in Scaffold v.5 (Proteome Software). To avoid double-counting mass spectra, exclusive spectral counts were used for the downstream proteomic analysis. Exclusive spectral counts were normalized using the normalized spectral abundance factor (NSAF) calculation^93^.

### 16S and 18S rRNA sequencing

RNA from one-fourth of each filter from incubation experiments was extracted from each filter piece with the Macherey-Nagel NucleoMag RNA kit on a Thermo Scientific Kingfisher Flex. cDNA was synthesized with the AzuraQuant™ cDNA synthesis kit according to the manufacturer’s instructions. Three amplicon libraries were then generated via a one-step PCR using the cDNA with the Azura TruFi DNA Polymerase PCR kit. The first library examined all three domains of life by simultaneously amplifying the V4-V5 region of the 16S and 18S rRNA genes with the 515F-Y (GTGYCAGCMGCCGCGGTAA) and 926R (CCGYCAATTYMTTTRAGTTT) primer set^94,95^. The second library specifically examined eukaryotes by amplifying the V9 region of the 18S rRNA gene with the 1389F (TTGTACACACCGCCC) and 1510R (CCTTCYGCAGGTTCACCTAC) primer set^96^. The third library also examined the V9 region of the 18S rRNA gene using cDNA from 3-51 µm and > 51 µm samples used for metatranscriptomics. The control treatment for the < 5 µm community was not included for sequencing with the 515F-Y and 926R primer.

Each reaction was performed with an initial denaturing step at 95°C for 1 minute followed by 30 cycles of 95°C for 15 seconds, 56°C for 15 seconds, and 72°C for 30 seconds. 2.5 µL of each PCR reaction was ran on a 1.8% agarose gel to confirm amplification. PCR products were purified using Beckman Coulter AMPure XP following the standard PCR clean-up protocol.

PCR quantification was performed using Invitrogen Quant-iT PicoGreen dsDNA Assay kit. Samples were then pooled in equal proportions followed by a 0.8x AMPure XP PCR purification. The final libraries were evaluated on an Agilent 2200 TapeStation (D1000 ScreenTape), quantified with Qubit HS dsDNA kit. The library with the 515F-Y/926R primers was sequenced on an Illumina NextSeq 2000 with a P1 flow cell (PE 300), and the library with the 1389F/1510R primers was sequenced on an Illumina NovaSeq X Plus with a 10B flow cell (PE 300). Mock communities were included in both libraries^97^.

Amplicons from both libraries were analyzed with QIIME2 v2024.5. Briefly, demultiplexed paired-end reads were trimmed to remove adapter and primer sequences with cutadapt^98^. Trimmed reads were then denoised with DADA2 to produce amplicon sequence variants (ASVs)^99^. The 515F-Y/926R library was analyzed to capture both 16S and 18S sequences as described in McNichol, et al. ^95^. Specifically, 16S and 18S sequences were split using bbsplit with 16S and 18S rRNA databases, 18S sequences were concatenated prior to DADA2, and then read counts were corrected for bias and random variation based on the relative abundances of 16S and 18S rRNA determined from both the TapeStation and DADA2 output. Taxonomic annotation of all ASVs was conducted with the q2-feature-classifier classify-sklearn naïve-bayes classifier^100,101^ and the SILVA database (Release 138.1) for 16S rRNA^102^ or the PR^2^ database (v5.1.0) for 18S rRNA^103^. All ASVs annotated as chloroplasts, mitochondria, or nucleomorphs were removed from the analysis. A large proportion of reads in the < 5 µm community were attributed to a single ASV annotated as the warm water diatom, *Hemiaulus senensis*, which we believe may be a mis-annotation in the PR^2^ database. Based on BLASTN results to NCBI’s core nucleotide database, this ASV was reannotated as *Navicula sp*.

### Cobalamin-related genes in prokaryotic genomes

NCBI accession numbers for genomes belonging to the genus *Polaribacter*, the family Nitrincolaceae, and the SAR92 clade were downloaded from GTDB (Data S2). When available, protein sequences for each genome were then downloaded with NCBI Datasets. Cobalamin synthesis (KEGG modules M00925 and M00122) and methionine synthase genes were queried using hmmsearch within KofamKOALA^86^. E-value cutoffs of 1 x 10^-5^ were used. Only genomes with greater than 80% completeness (CheckM2) were used.

### Statistical analyses

Two-way ANOVAs followed by Tukey’s honest significant difference test were performed on the biological and chemical seawater measurements using Graphpad PRISM v10.5.0.

## Data availability

An overview of the publicly available data from the CICLOPS expedition is on Zenodo: https://doi.org/10.5281/zenodo.1728790. Hydrographic and nutrient data are deposited in BCO-DMO: https://www.bco-dmo.org/project/774945. Raw metabarcoding and metatranscriptomics sequences are deposited in the NCBI Sequence Read Archive under BioProjects PRJNA1499736 and PRJNA890306 respectively. The metatranscriptome assembly is deposited in Zenodo: https://doi.org/10.5281/zenodo.17307449. The mass spectrometry proteomics data have been deposited to the ProteomeXchange Consortium via the PRIDE partner repository under the identifiers PXD084199 and PXD084282.

## Acknowledgements

We thank the captain and crew of the RVIB *Nathaniel B. Palmer.* We also thank Paloma Lopez for assistance with the protein extractions, and Adrian Marchetti for comments.

## Author Contributions

Conceptualization: G.R.D. and M.A.S., Data Curation: R.H.L., D.R., R.M.K., R.J.C., F.B., R.C., C.B., and G.R.D., Formal analysis: R.H.L., R.C., C.B., G.R.D., and M.A.S., Funding acquisition: G.R.D. and M.A.S., Investigation: R.H.L., D.R., R.M.K., R.J.C., N.L.S., S.K., L.E.L., F.B., O.M., R.C., C.B., H.Z., D.M.M., A.E.A., M.R.M., G.R.D., and M.A.S. Project administration: G.R.D. and M.A.S., Resources: O.M., R.C., C.B., A.E.A., G.R.D., and M.A.S., Supervision: G.R.D. and M.A.S., Visualization: R.H.L., Writing — original draft: R.H.L. and M.A.S., Writing — review & editing: R.M.K, S.K., L.E.L., R.C., C.B., and G.R.D.

## Competing interests

The authors declare no competing interests.

Supplementary Information is available for this paper.

Correspondence and requests for materials should be addressed to Mak Saito.

