## Supplementary Information for "Primary vitamin B_12_ limitation and widespread stress in Antarctic phytoplankton"

**Figure S1.** Additional measurements from the incubation experiment: (A) Photosynthetic efficiency ($F_v/F_m$), (B) Biogenic silica (µmol $L^{-1}$), (C) Phosphate (µmol $L^{-1}$), (D) Silicic acid (µmol $L^{-1}$). For Fv/Fm and biogenic silica, error bars represent the standard deviation of the mean (*n* = 3).

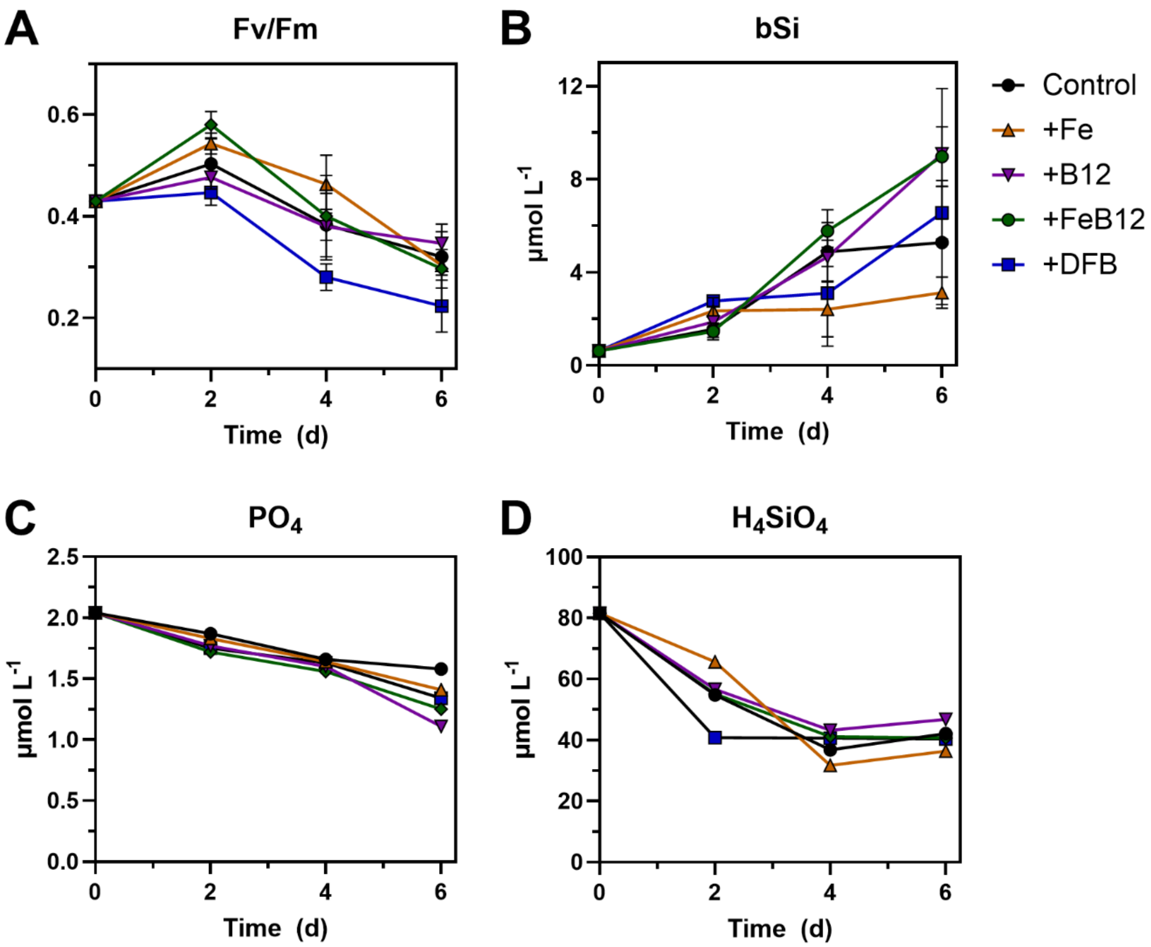

**Figure S2.** (A) Cobalamin uptake (%) and (B) cobalt uptake rates (pmol $L^{-1}$ $d^{-1}$) at Stations 4, 11, and 15 for the ≥ 0.2 µm community (solid) and ≥ 3.0 µm community (dashed) (C) Total (solid) and labile (dashed) dissolved cobalt concentrations. Dissolved cobalt data are reproduced from Chmiel et al. (*1*).

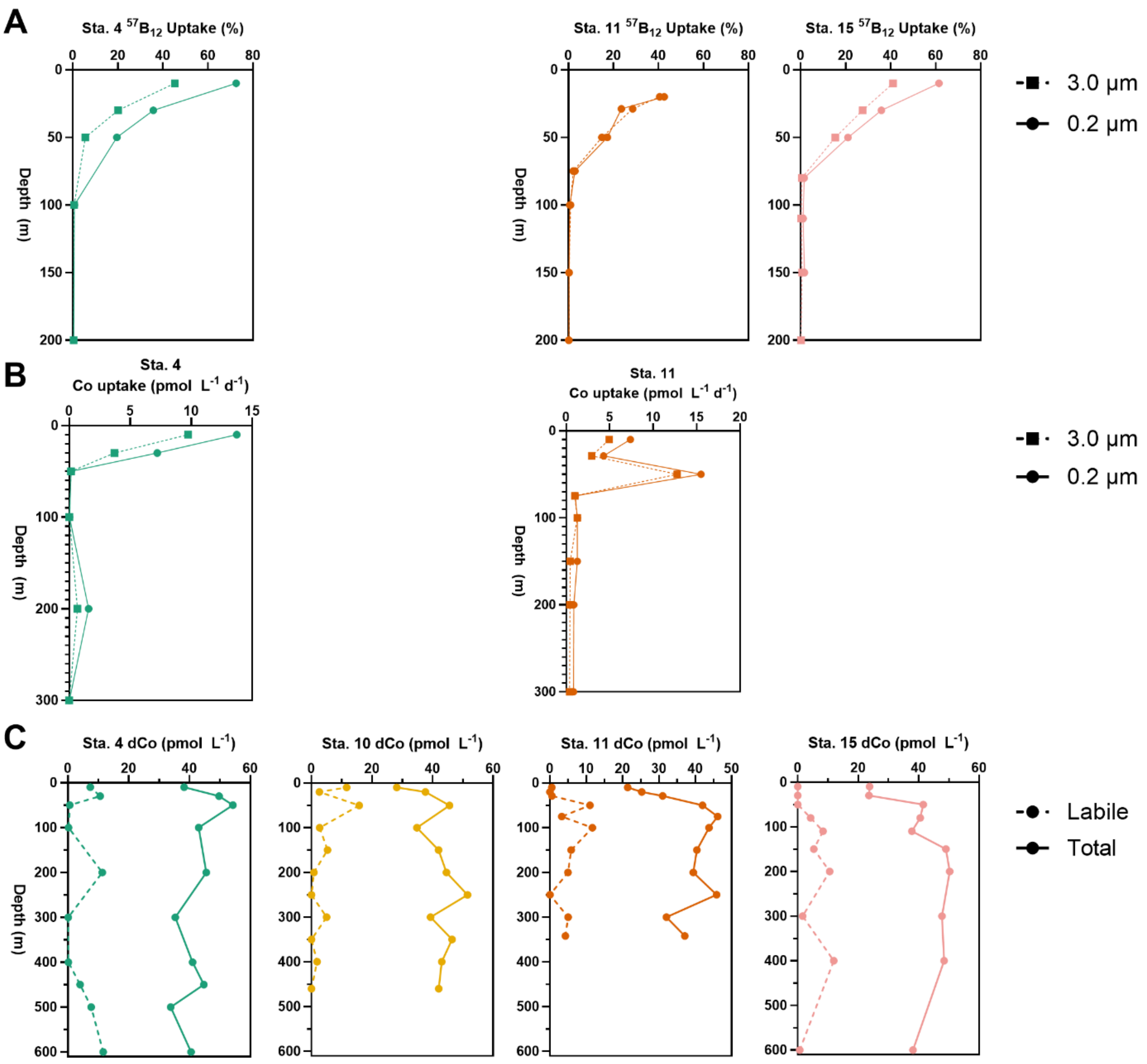

**Figure S3.** Temperature (ºC) versus salinity at the experimental site (Station 4). Endmembers are labeled as modified Circumpolar Deep Water (mCDW), Winter Water (WW), and Antarctic Surface Water (AASW) (*2*).

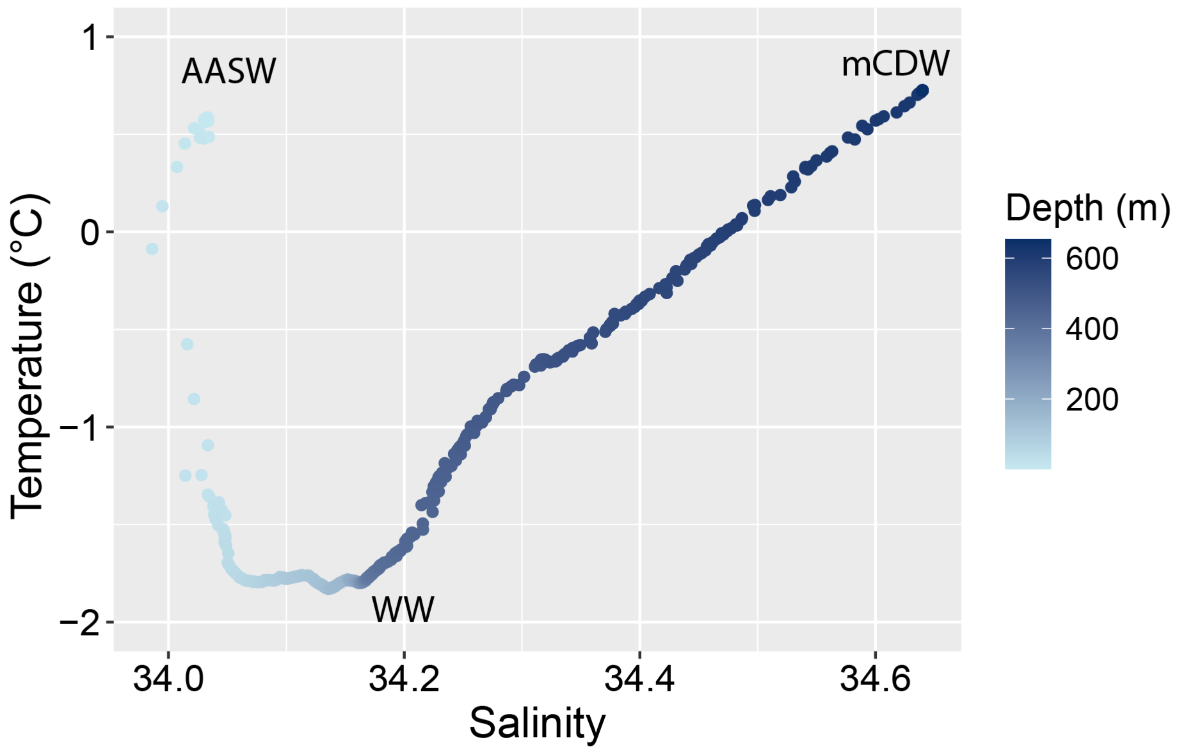

**Figure S4.** Vertical profiles of copper (Cu), nickel (Ni), and cadmium (Cd) concentrations (nmol $L^{-1}$) among stations.

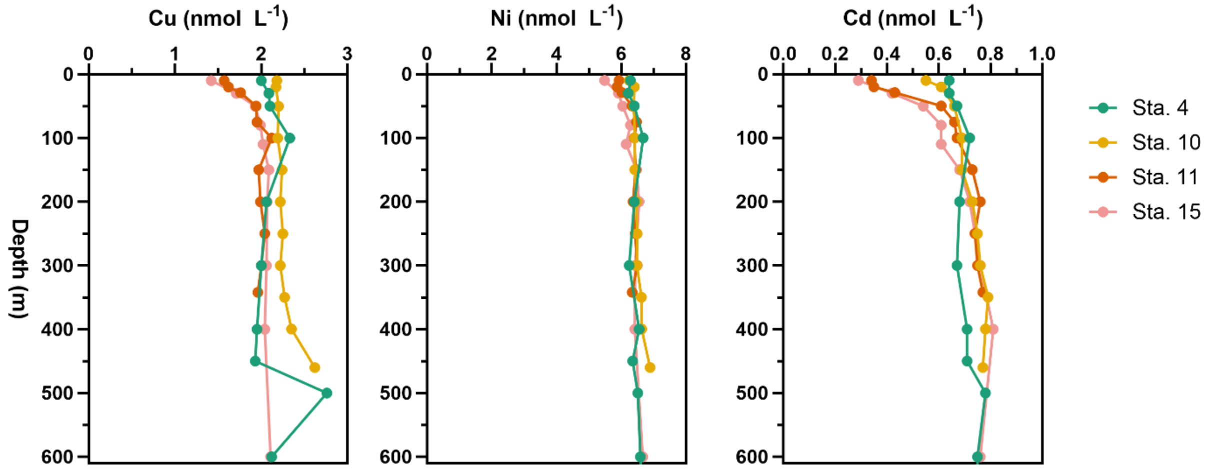

**Figure S5.** Relative abundances of diatoms versus *Phaeocystis* in metaproteomic samples from surface seawater. Station 4 is denoted with a + symbol.

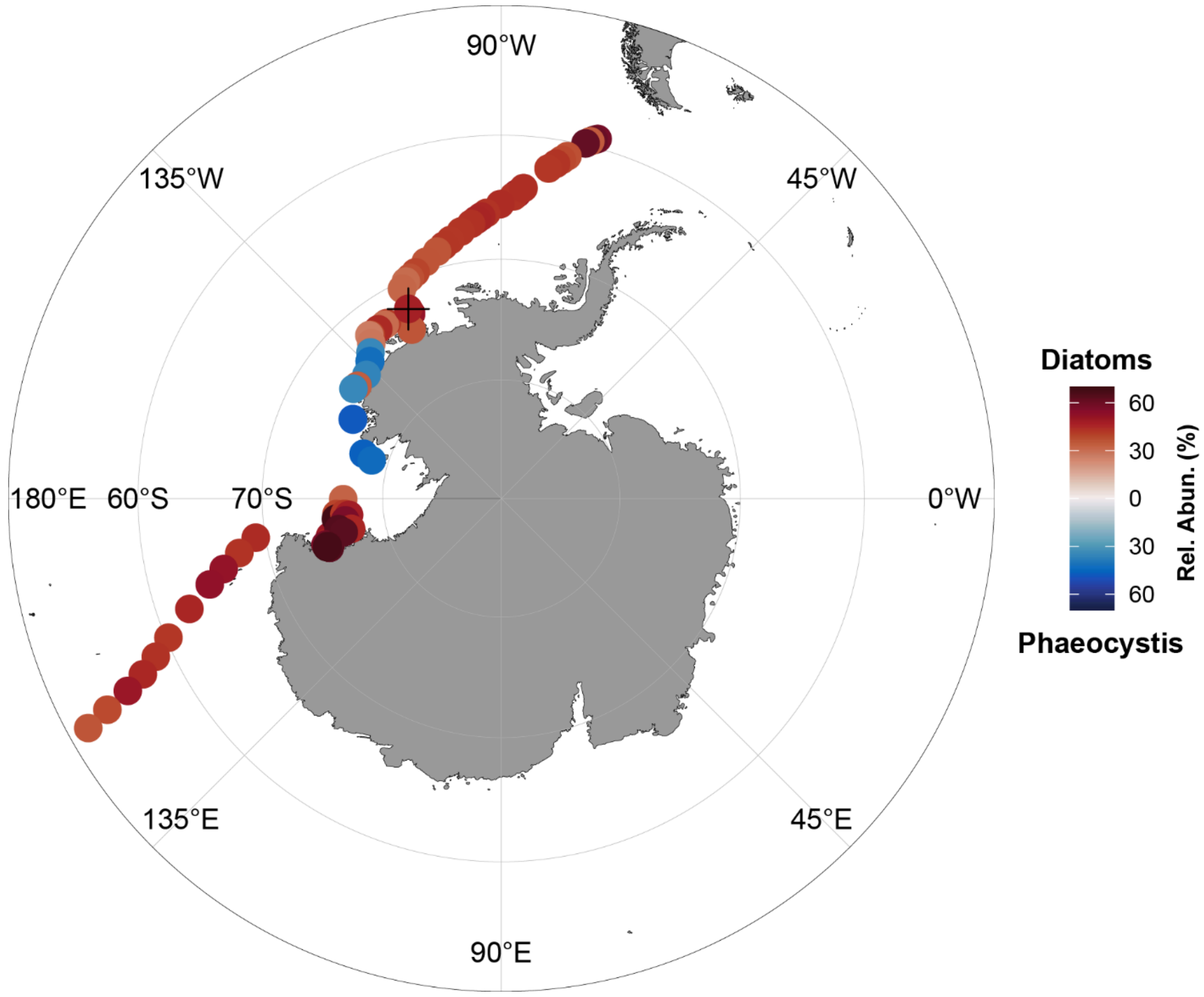

**Figure S6.** Additional pigment concentrations (ng $L^{-1}$) from the incubation experiment: (A) 19'-hexanoyloxyfucoxanthin, (B) lutein, and (C) zeaxanthin. Error bars represent the standard deviation of the mean ($n$ = 3).

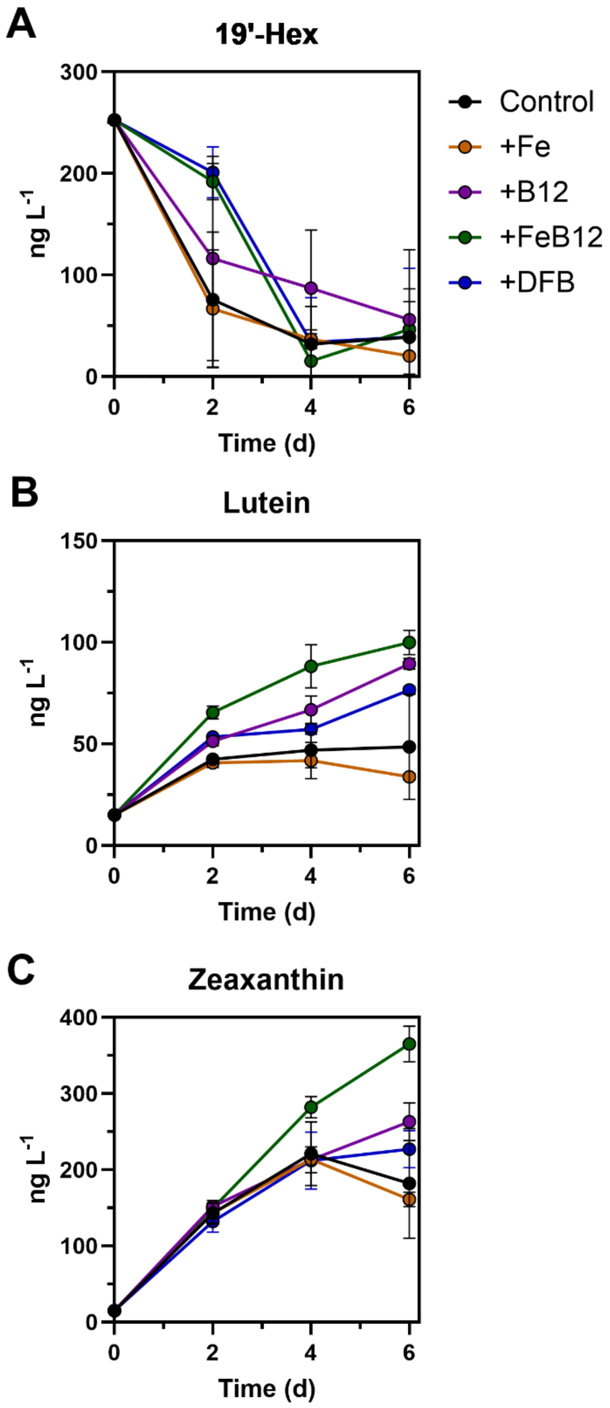

**Figure S7.** Relative abundances of diatom genera from 18S rRNA within each size fraction and treatment.

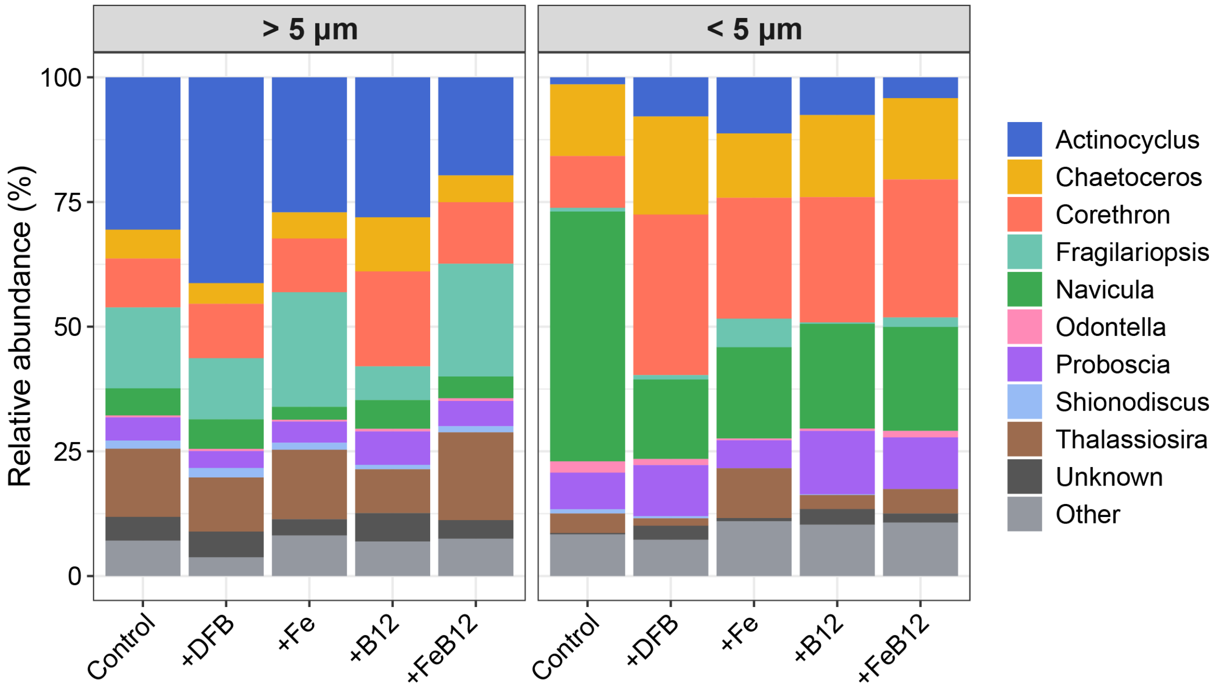

**Figure S8.** Relative abundances of eukaryotic lineages from 18S rRNA at Stations 4, 11, 15, and 27. Samples were collected at multiple depths at each station and size fractionated to capture the 3.0-51 µm (left) and > 51 µm (right) communities.

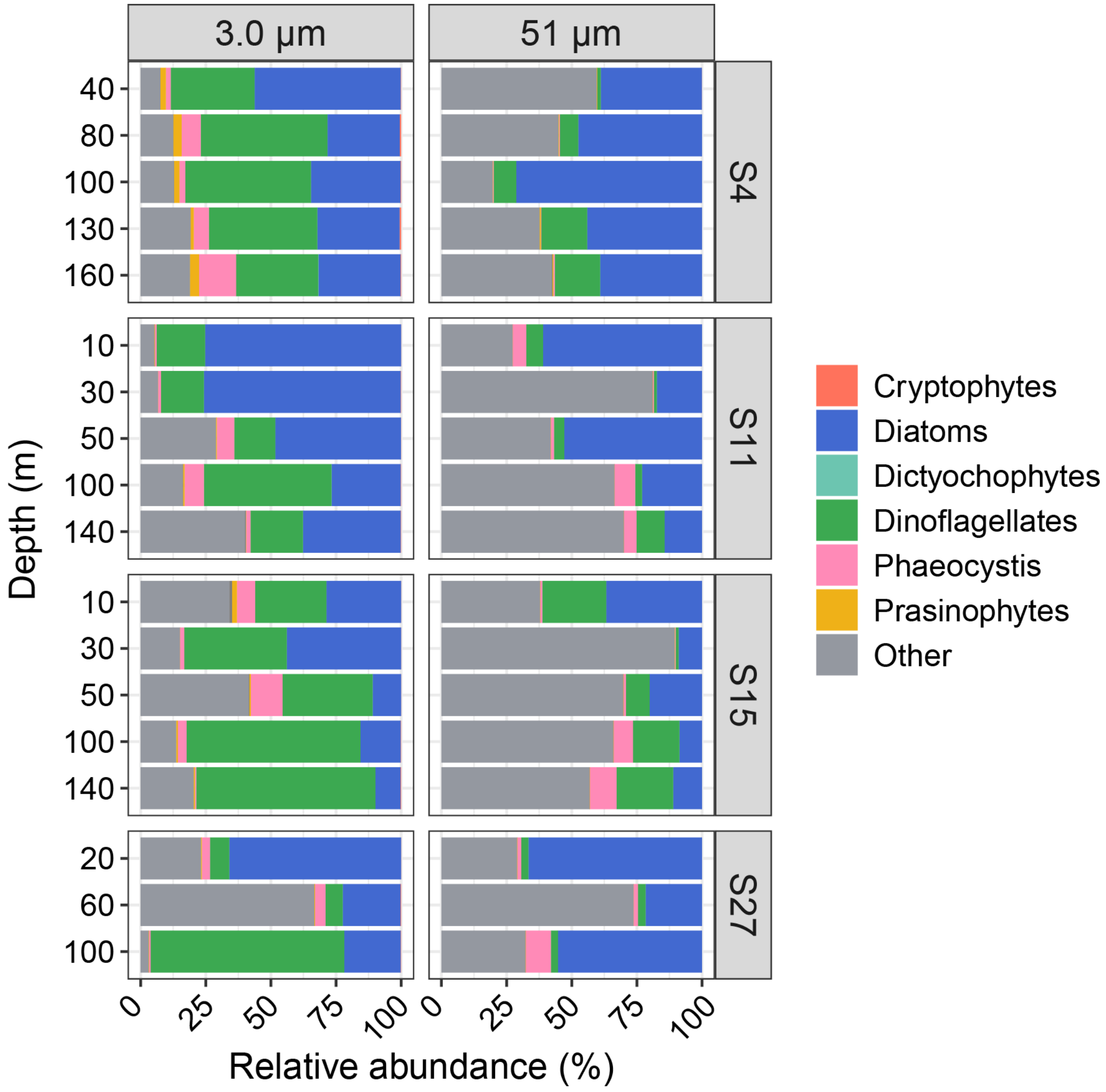

**Figure S9.** Relative abundances of diatom genera within the diatom community at Station 4. Samples were collected at multiple depths and size fractionated to capture the 3.0-51 µm (left) and > 51 µm communities (right).

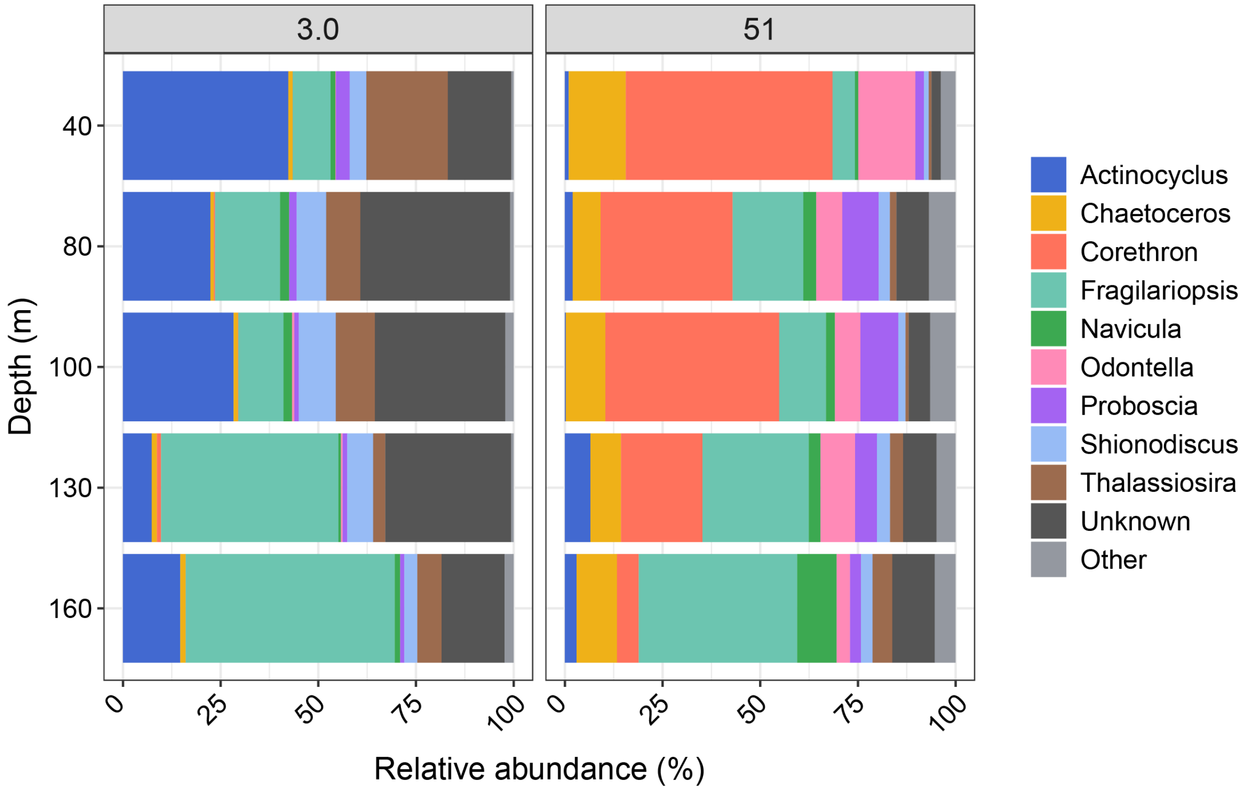

**Figure S10.** Prokaryotic abundances from flow cytometry (cells $mL^{-1}$). (A) Abundances over time in each of the experimental treatments. (B) Comparison with previous studies of relative increases in chlorophyll *a* concentrations with +B12 or +FeB12 treatments compared to +Fe treatments versus prokaryotic abundances (*3, 4*).

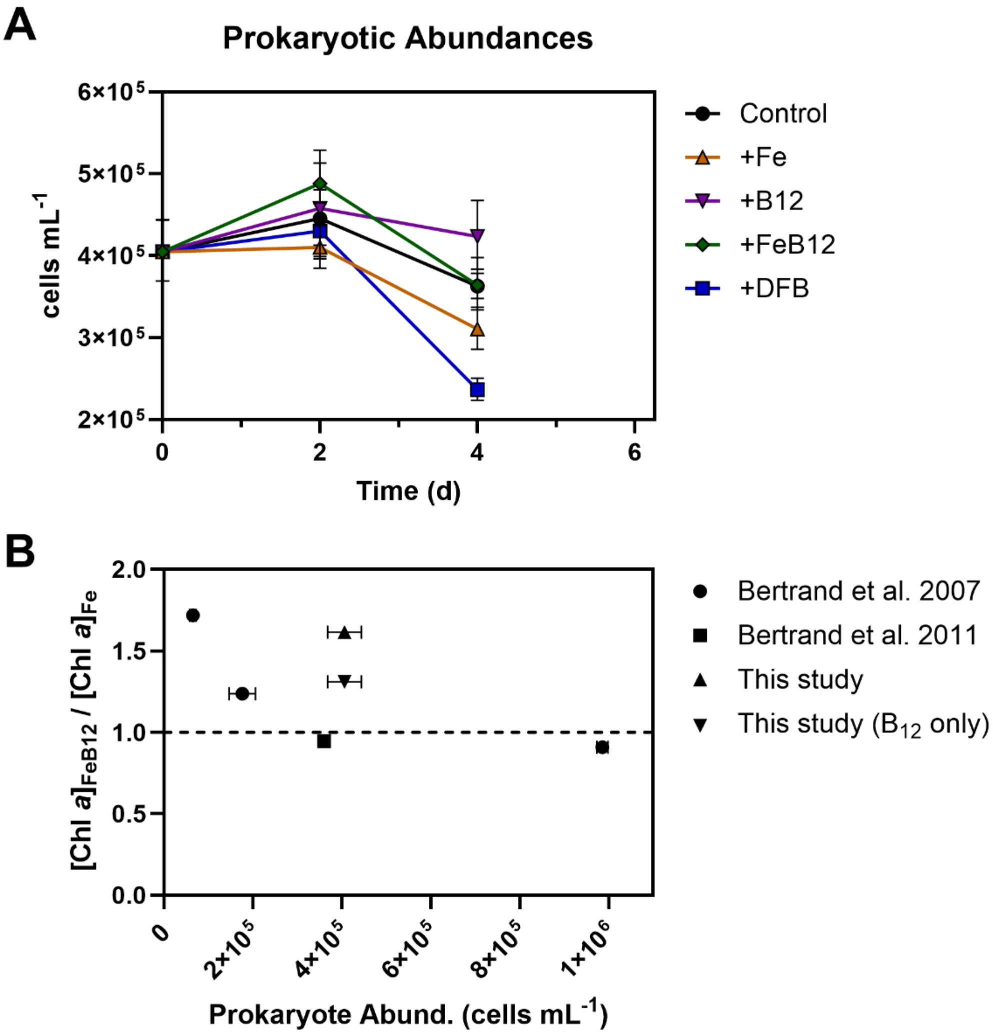

**Figure S11.** Relative abundances of proteins from major taxonomic groups at Stations 4, 11, 15, and 27. Samples were collected at multiple depths at each station and size fractionated to capture the 0.2-3.0 µm (left), 3.0-51 µm (middle) and > 51 µm (right) communities.

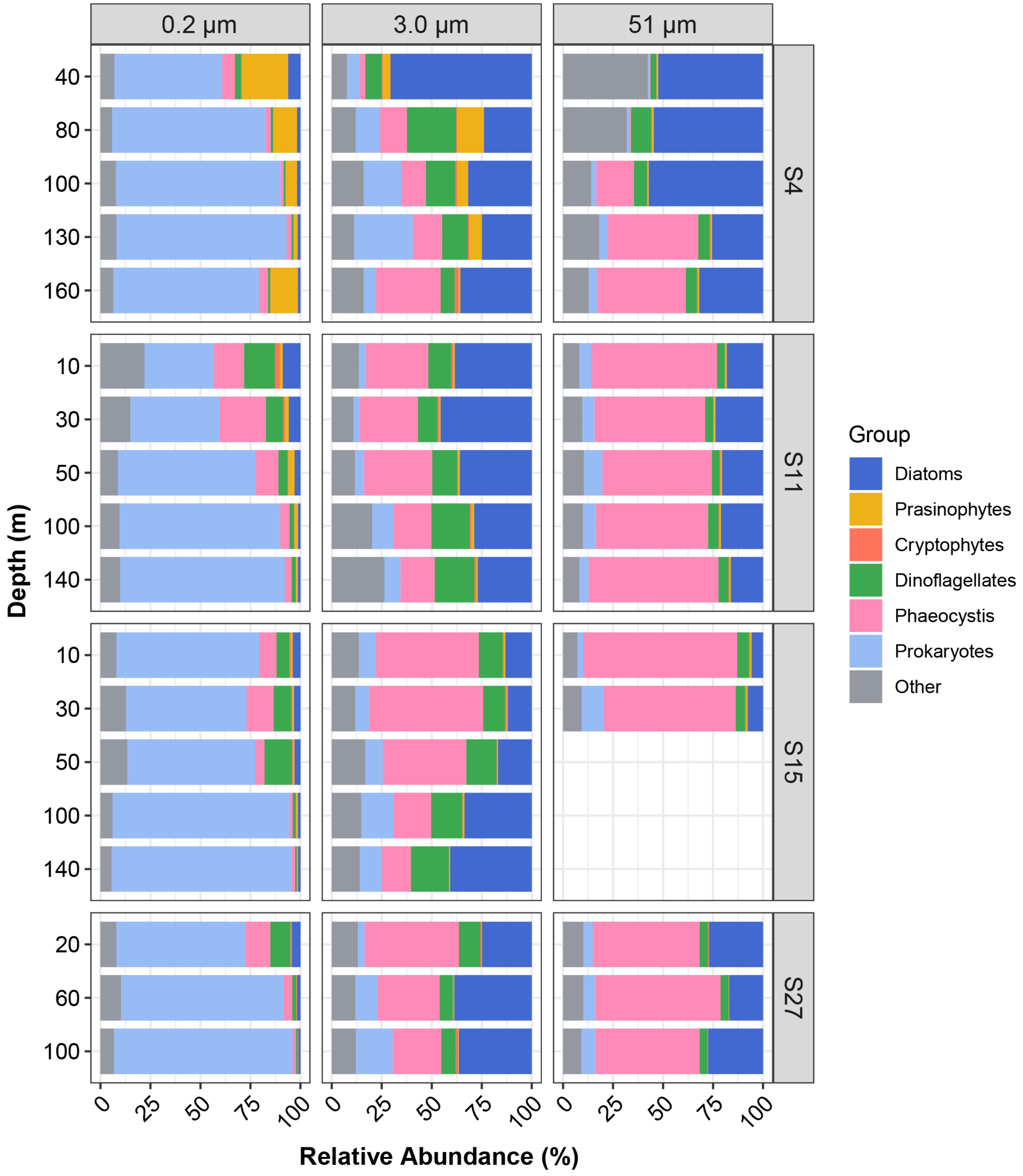

**Figure S12.** Prokaryotic genomic content for cobalamin and methionine synthesis in Nitrincolaceae (*n* = 623), *Polaribacter* (*n* = 644), and SAR92 clade bacteria (*n* = 10). (A) Percent of genomes with each cobalamin synthesis gene. (B) Cobalamin synthesis pathway completeness. (C) Percent of genomes with metH, metE, both, or neither.

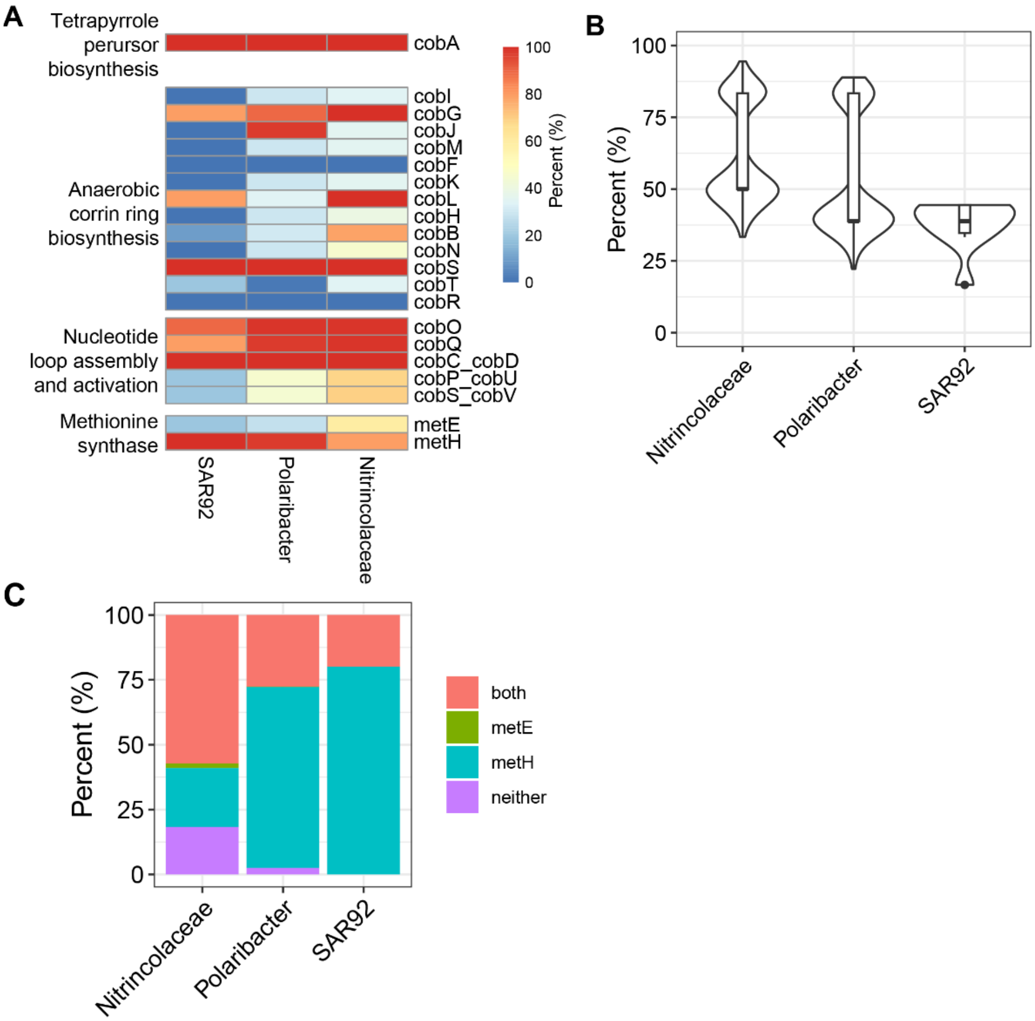

**Figure S13.** Percent CBA1 abundances within diatoms from underway samples on a $\log_{10}$ scale.

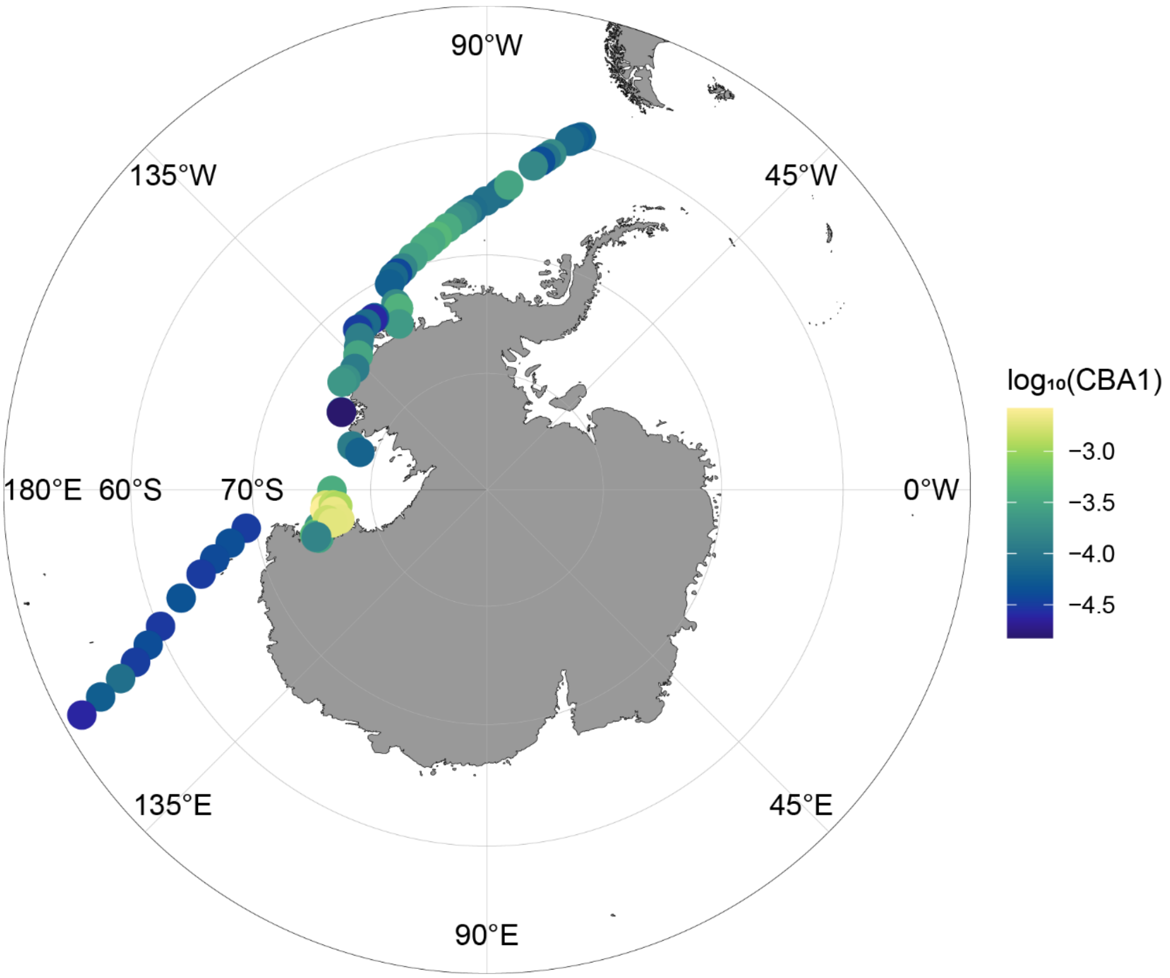

**Figure S14.** Sea ice concentrations in the Amundsen Sea from November 30, 2017 with stations denoted.

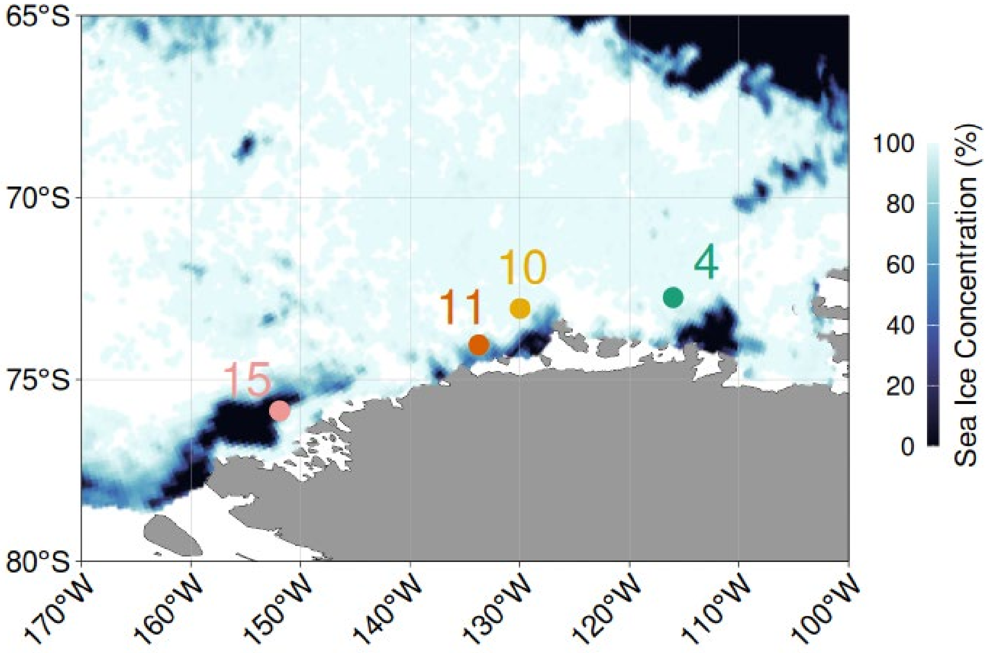

**Table S1.** Taxon-specific responses to nutrient addition in Antarctic polynyas under primary cobalamin and secondary iron limitation.

| | +Fe | $+B_{12}$ | $+FeB_{12}$ |
|---|---|---|---|
| **Diatoms** | **-** | + | ++ |
| ***Phaeocystis*** | **-** | **-** | **-** |
| **Prasinophytes** | **-** | + | ++ |
| **Cryptophytes** | + | **-** | **-** |

**Data S1. (separate file)** Underway sample metadata.

**Data S2. (separate file)** Accession numbers and taxonomy for the bacteria genomes used in Figure S12.
